# A microeukaryotic PPR-DYW protein supports multisite C- and A-deamination RNA editing

**DOI:** 10.64898/2026.09.04.749523

**Authors:** Matus Valach, Lisbeth C. Aguilar, Ninaad Kalla, Poorya Mirzavand Borujeni, B. Franz Lang, Marlene Oeffinger, John M. Pascal, Reza Salavati, Gertraud Burger

## Abstract

RNA editing in mitochondria is vital for many eukaryotes. In plants, mitochondrial C-to-U deamination editing is catalyzed by tens to hundreds of PPR-DYW proteins, each typically dedicated to a specific site. Where retained, PPR-DYW family expanded independently in all eukaryotic groups examined here—except in marine microeukaryotes diplonemids, which encode a single homolog despite deaminating their mitochondrial RNA at more than 100 sites. Here we characterize this unconventional deaminase, PPRD1, from *Diplonema papillatum*. Native affinity pulldown of the protein identified 20 predominantly sub-stoichiometric partners, including potential RNA-binding helical-repeat proteins. Among them, the divergent PolX-like protein DAPX1 consistently and reciprocally co-purified with PPRD1 in near-equal proportions. Structural modelling suggests that DAPX1 may stabilize the deaminase catalytic domain and expand its interaction interface. Silencing either PPRD1 or DAPX1 inhibited cell growth and reduced *in vivo* not only C-to-U, but also A-to-I deamination across five mitochondrial RNA-editing clusters encompassing 110 sites. Together, these results support a model of PPRD1 and DAPX1 forming the core of the *Diplonema* deamination-editing machinery, with sub-stoichiometric partners acting as specificity factors that guide accurate RNA editing with minimal off-target effects.

## INTRODUCTION

RNA base-modifying enzymes play an important role in gene expression by influencing transcript stability, coding potential, and translation efficiency [1]. Among these, RNA deaminases form a major class: adenosine deaminases acting on RNA (ADARs) catalyze adenosine-to-inosine (A-to-I) editing of double-stranded RNAs in animals, altering codons and splicing patterns [2]; cytidine deaminases such as APOBEC1 mediate cytidine-to-uridine (C-to-U) editing in mammalian mRNAs [3]; and tRNA adenosine and cytidine deaminases (ADATs, CDATs) modify tRNA bases in the nucleus and mitochondria [4].

A particularly intriguing class of RNA deaminases is involved in organelle RNA editing. In plant organelles, site-specific C-to-U editing, predominantly in mRNAs, frequently restores conserved codons. The well-characterized enzymes mediating this editing are PPR-DYW proteins, which comprise an N-terminal pentatricopeptide repeat (PPR) array and a DYW catalytic domain belonging to the cytidine deaminase superfamily (reviewed in [5]). Biochemical reconstitution [6,7] and heterologous expression assays [8–10] established the DYW domain as the catalytic deaminase and showed that complete PPR-DYW proteins can perform site-specific editing without additional plant factors. The PPR array confers target specificity by recognizing an RNA sequence motif and positioning the deaminase domain at the cognate cytidine approximately four nucleotides downstream. This recognition-and-positioning mechanism has recently been visualized in an RNA-bound PPR-DYW structure [11]. In other plant editing systems, RNA recognition and catalysis are split between a PPR protein lacking a complete DYW domain and a separate DYW protein recruited in trans (reviewed in [12,13]).

PPR-DYW-like proteins have also been reported outside land plants, first in the heterolobosean *Naegleria* and subsequently in other protists [14,15]; however, the biochemical functions of these proteins in protists remain uncharacterized. At the same time, mitochondrial transcriptome analyses across several protist lineages have revealed substitutions consistent with deaminase-mediated RNA editing, but both the underlying mechanism and the proteins responsible for these reactions are still unknown [16–18]. Diplonemids, a group of marine microeukaryotes with unusually complex mitochondrial gene expression [19] (**Figure 1A**), provide a tractable system for investigating the molecular basis of such editing. In the diplonemid type species *Diplonema papillatum* (hereafter *Diplonema*), C-to-U and A-to-I substitutions occur at highly specific sites, predominantly within defined, dense clusters. Importantly, inosine has been demonstrated directly at A-to-I editing sites in both mitochondrial mRNA and rRNA, providing experimental evidence that these substitutions arise by deamination rather than being inferred solely from genomic-to-transcript sequence differences [20,21].

**Figure 1.**
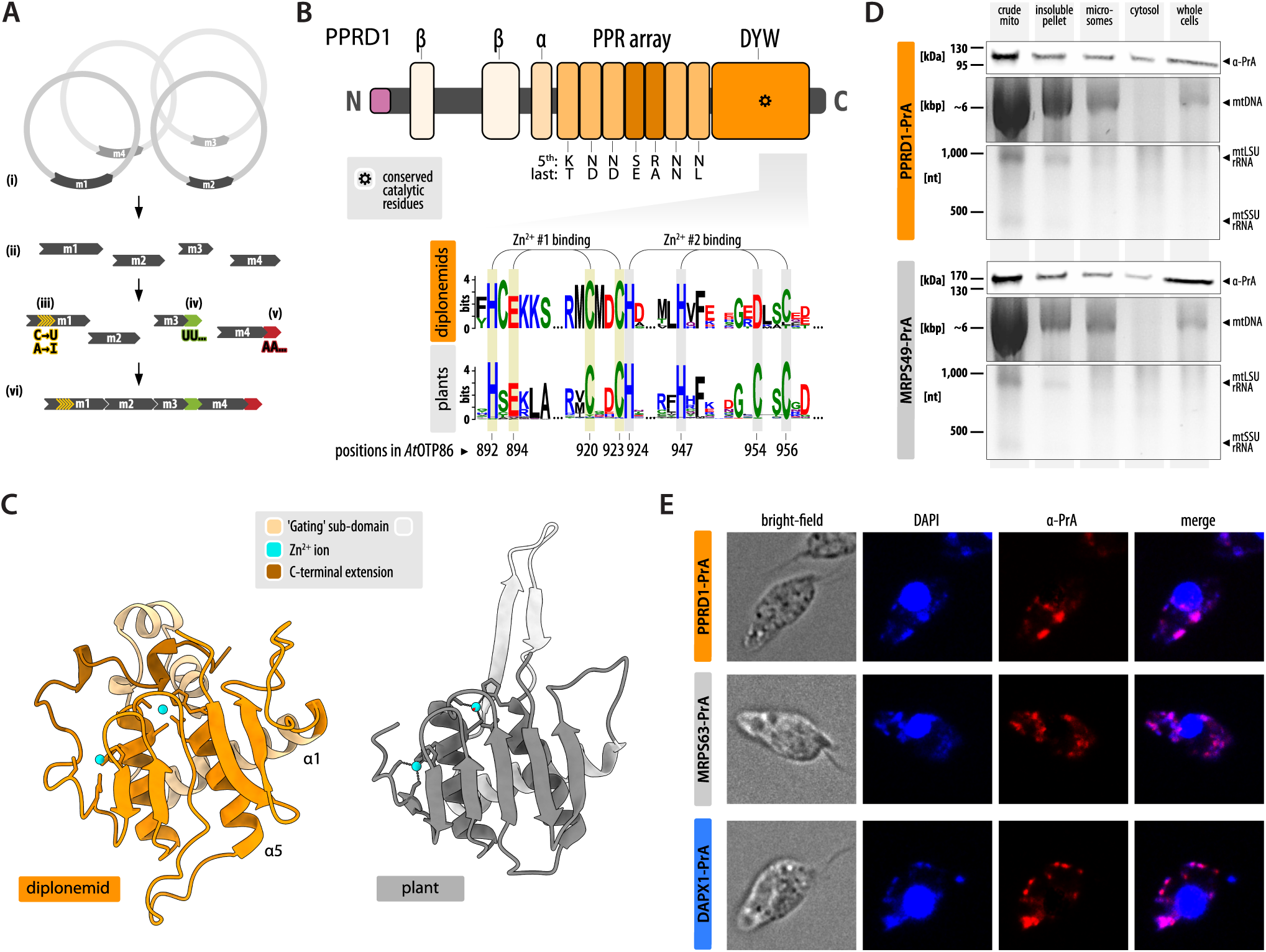
Diplonemid mitochondrial gene expression involves deamination-based RNA editing, and PPRD1 is a mitochondrial protein displaying features consistent with this activity. (**A**) Diplonemid mitochondria contain 30 to nearly 200 distinct chromosomes, each typically encoding a short gene fragment termed ‘module’ (i). After transcription into separate short RNAs (ii), certain modules undergo site-specific RNA editing by deamination (iii) and 3′ uridylation (iv), followed by polyadenylation of terminal fragments (v). The processed modules are then joined to form a contiguous mature transcript (vi). (**B**) Structural features of *D. papillatum* PPRD1 include two β-sheet bundles, a coiled-coil α-helix, a pentatricopeptide repeat (PPR) array, and a C-terminal DYW catalytic domain (shown in orange shades). The mitochondrial targeting pre-sequence (MTP) is highlighted in light purple. The PPR array contains 35– and 31-residue motifs (lighter and darker shades, respectively). Residues at the fifth and last positions of PPR motifs can mediate RNA base recognition [29,30]. In diplonemid PPRD1, these residues only occasionally conform to canonical combinations, notably N–D, N–N, and N–L, which show preference for pyrimidines. Sequence logos derived from DYW domain alignments show conserved zinc-binding and catalytic residues characteristic of plant DYW deaminases; residue-numbering was based on *Arabidopsis thaliana* OTP86. (**C**) Comparison of the spatial arrangement of key structural features (active-site residue, zinc-binding residues, and proposed regulatory switch) in the predicted *Diplonema* PPRD1 DYW domain and the experimentally determined *At*OTP86 structure. Regions corresponding to the flexible ‘gating’ sub-domain are shown in lighter shades [7,32] (see also **Supplementary Figure S3**) (**D**) Distribution of PrA-tagged PPRD1 and the mitoribosomal protein MRPS49 across subcellular fractions separated by sucrose-gradient velocity sedimentation, assessed by Western blotting. Mitochondrial DNA and ribosomal RNA separated by agarose gel electrophoresis serve as fractionation markers (see also **Supplementary Figure S5**). (**E**) Immunofluorescence microscopy showing subcellular localization of PrA-tagged PPRD1, the mitoribosomal protein MRPS63, and DAPX1, visualized using anti-PrA antibodies and DAPI-based DNA staining.

A recent *in silico* study of the *Diplonema* mitochondrial proteome identified two candidate nucleus-encoded RNA deaminases [22]. DIPPA_33495 is a CDAT homolog, which in trypanosomes converts C to U at the wobble position of the anticodon, thereby allowing the nucleus-encoded, imported tRNA-Trp to decode mitochondrial UGA codons as tryptophan [23,24]. The second candidate, DIPPA_21441, designated PPRD1 (PPR-containing Deaminase 1), resembles plant PPR-DYW proteins in containing a PPR array and a DYW-like domain [22].

PPRD1 has two unusual features that distinguish the *Diplonema* homolog from characterized plant PPR-DYW proteins. First, plants have tens to hundreds of PPR-DYWs, with individual proteins typically associated with one or a few editing sites, whereas *Diplonema* has only a single homolog despite its 114 mitochondrial deamination sites. Second, characterized PPR-DYW proteins catalyze C-to-U editing, whereas editing in *Diplonema* comprise both C-to-U (85 sites) and A-to-I (29 sites) substitutions [25,26]. These observations raise two central questions: whether PPRD1 catalyzes both reactions and, if so, how a single PPR-DYW protein specifically recognizes numerous editing sites. To address these questions, we characterized the structural and functional properties of PPRD1 from *Diplonema*, defined the network of associated proteins, and tested the roles of PPRD1 and its major partners in mitochondrial deamination editing.

## RESULTS

### 1. PPRD1 combines an atypical PPR array with a structurally canonical DYW domain

PPRD1 was identified in all 12 analyzed diplonemid species, with the inferred proteins sharing ∼60% sequence identity (**Supplementary Figure S1**). The protein contains a central tandem array of seven PPR motifs (**Figure 1B**), which, despite their size variability, all belong to the P type [22], as is typical of non-plant lineages [27] (but see, e.g., [28] and **Supplementary Information** for more details). Because two residues at the fifth and last positions in each PPR-motif collectively determine base specificity [29,30], an array of seven motifs can theoretically recognize a seven-nucleotide RNA sequence. However, in diplonemid PPRD1 homologs, these residues only sporadically match canonical base-recognition combinations as defined in the plant PPR code. Even when they do (**Figure 1B**), the pairs (e.g., N-D, N-N, N-L) predominantly specify pyrimidines and together provide limited information for defining a unique RNA target. Thus, according to the canonical PPR recognition code, the PPRD1 residue pairs do not readily explain specific recognition of the numerous mitochondrial editing sites in *Diplonema*.

The C-terminal region of PPRD1 harbours a 120-residue-long DYW domain (IPR032867/PF14432) that has all the hallmarks of cytidine deaminases, including the conserved residues coordinating the catalytic and structural Zn2+ ions [22] (**Figure 1B**). Computational protein structure prediction yielded an overall high-confidence model for PPRD1, with lower score regions restricted to the termini and inter-domain loops (**Supplementary Figure S2A**). The *Diplonema* DYW domain closely conformed to experimentally determined *Arabidopsis* structures, with differences largely confined to the flexible ‘gating ‘subdomain (**Figure 1C, Supplementary Figure S3**; for more details, see **Supplementary Information**). Stereochemistry-based prediction of zinc-binding sites [31] identified two well supported sites in PPRD1 corresponding to those in *Arabidopsis* DYW structures [7] (**Supplementary Figure S3**). Together, these results support the classification of PPRD1 as a DYW-family deaminase with an atypical PPR array.

### 2. A single DYW deaminase in diplonemids contrasts with family expansions elsewhere

The presence of only a single PPR-DYW protein per diplonemid, despite extensive mtRNA deamination editing, prompted us to survey PPR-DYW family diversity across Discoba and malawimonads. The latter also encode PPR-DYW proteins [15] and were included as a phylogenetically informative deep-branching and slowly evolving counterpart to Discoba [33]. Searches of available proteomes identified multiple PPR-DYW paralogs in *Malawimonas californiana*, *Gefionella okellyi*, and *Acrasis kona*, but none in *M. jakobiformis*, *Andalucia godoyi*, *Euglena gracilis*, or *Trypanosoma brucei*. Phylogenetic analysis was consistent with lineage-specific expansions in malawimonads and heteroloboseans and losses in several other lineages (**Supplementary Figure S4**, **Supplementary Information**). Similar lineage-specific expansions have been well documented in plants [34], further highlighting the distinctive single-copy state of PPRD1 in diplonemids.

### 3. PPRD1 is localized to mitochondria

We next tested whether PPRD1 has the mitochondrial localization expected for a factor implicated in mitochondrial RNA editing and predicted computationally [22], using a *Diplonema* cell line that expresses PPRD1 fused to a Protein A (PrA) tag. (Primers, constructs, and tagged cell lines used in this study are listed in **Supplementary Table S1–S3**.) Following separation of the cell lysate on a discontinuous sucrose velocity gradient, tagged PPRD1 showed a distribution resembling that of the mitoribosomal protein MRPS49 [35], mitochondrial DNA (mtDNA) and mt-rRNAs (**Figure 1D; Supplementary Figure S5**). Mitochondrial localization of the fusion protein was further supported by anti-PrA immunofluorescence microscopy and co-staining with the DNA-binding dye DAPI. The tagged PPRD1 co-localized with mtDNA, similar to the pattern observed for another confirmed mitochondrial protein, the mitoribosomal protein MRPS63 [35] (**Figure 1E**).

### 4. Affinity purification identifies the PPRD1–DAPX1 core associating with diverse substoichiometric partners

We tested whether PPRD1 is part of a larger protein assembly using single-step affinity purification. PPRD1-PrA and co-purifying proteins were captured from a cell lysate on IgG-coated beads [35,36]. Liquid chromatography–tandem mass spectrometry (LC-MS/MS) of multiple biological replicate pulldowns identified, in addition to PPRD1, 17 high-confidence protein associates (**Figure 2A**, left).

**Figure 2.**
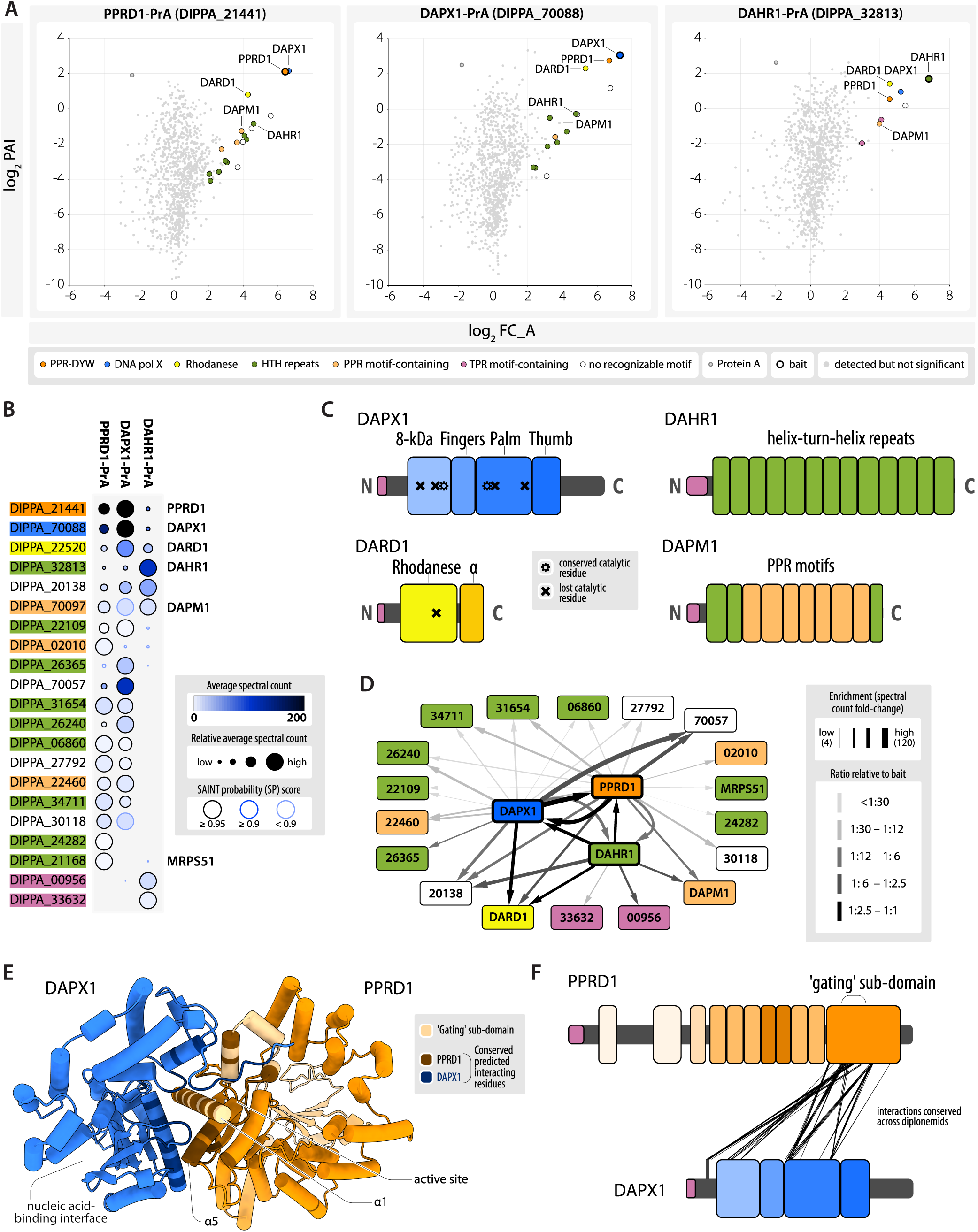
PPRD1 together with its partner DAPX1 forms the core of the putative mitochondrial RNA deaminase complex, acting as the central hub in a multi-protein network. (**A**) Enrichment-vs-abundance plots of proteins co-purifying with PrA-tagged PPRD1 (DIPPA_21441, left panel), DAPX1 (DIPPA_70088, middle), and DAHR1 (DIPPA_32813, right). All affinity purification and mass spectrometry experiments were performed at least in triplicate. Proteins were classified based on their abundance (PAI), enrichment (average fold change; FC_A), and probability score (SP) (for details, see **Supplementary Table S5**). Significant proteins (FC_A > 4, SP > 0.9) were coloured in line with their characteristic domains or sequence motifs (e.g., DNA PolX, PPR, TPR). DAPM1 was detected in DAPX1-PrA pulldowns (middle panel, labeled), but fell just below the significance threshold for prey inclusion.(**B**) Dot plot of relative protein abundances across affinity purification experiments with the three different baits, measured in average spectral counts (see the key for details). Protein IDs are highlighted in the same categorization hues as in (**A**) and clustered based on similar abundance and distribution across the three pulldowns. (**C**) Structural and functional features and domains of the four most significant partners of PPRD1. (**D**) Combined interactome of PPRD1, DAPX1, and DAHR1. Nodes represent detected proteins and edges indicate significant associations with the respective baits. Protein IDs are highlighted in the same categorization hues as in (**A**). (**E**) Predicted structure of the PPRD1–DAPX1 binary complex. Interacting residues (shown in darker shades; see also **Supplementary Figure S7**) were defined based on residue pair distances (≤4 Å) in the predicted complex structure and restricted to interactions conserved across diplonemids within regions of reliable local structure (**Supplementary Figure S8**). The interface involves predominantly the α1 helix of the ‘gating ‘sub-domain and the C-terminal end of the α5 helix carrying the catalytic residue of PPRD1. The relative orientation of the two subunits leaves the nucleic acid-binding side of the PolX domain and the deaminase active site accessible, albeit the PPR array partially approaches the latter in the predicted structure. Low-confidence regions of pLDDT<50 at N– and C-termini are not shown. (**F**) Locations of the predicted high-confidence interface residues along the sequences of PPRD1 and DAPX1. (For domain delimitations, see (**B**) and **Figure 1B**.)

Among the most prominent partners were DAPX1 (<u>D</u>eaminase <u>A</u>ssociated <u>P</u>olymerase <u>X</u>-like 1; DIPPA_70088) and DAHR1 (<u>D</u>eaminase <u>A</u>ssociated <u>H</u>elix–turn–helix <u>R</u>epeat 1; DIPPA_32813), both discussed in further detail below. These two proteins were used as PrA-tagged baits in analogous affinity-purification experiments (**Supplementary Table S4**). This resulted in the addition of three significant proteins, bringing the total to 20 high-confidence associates of PPRD1 (**Figure 2B**, **Supplementary Figure S6**). Bait-specific prey abundances, enrichments, interaction scores, localization predictions, and taxonomic distributions are compiled in **Supplementary Table S5**.

Several PPRD1 partners, all predicted to localize to mitochondria [22], are of particular interest. DAPX1 mentioned above contains a conserved DNA polymerase type X family domain (**Figure 2C**) including a region corresponding to the DNA polymerase β thumb subdomain (IPR029398/PF14791), which contributes to DNA binding in active PolX enzymes. However, most residues required for polymerase activity are substituted, arguing against canonical polymerase activity of DAPX1 (**Supplementary Figure S7**). Outside the PolX domain, DAPX1 lacks significant sequence similarity to characterized proteins from non-diplonemid organisms. Mitochondrial localization of DAPX1 was confirmed by immunofluorescence microscopy (**Figure 1E**). It co-purified with PPRD1 at an approximately 1:1 ratio when either PPRD1 or DAPX1 was used as bait, and the two proteins were again recovered at near-equimolar levels in DAHR1 pulldowns, consistent with DAPX1 and PPRD1 forming a core association.

Among the substoichiometric proteins co-purifying with both PPRD1 and DAPX1 (**Figure 2A**), DARD1 (<u>D</u>eaminase <u>A</u>ssociated <u>R</u>hodanese <u>D</u>omain 1; DIPPA_22520) was the most abundant. In PPRD1 and DAPX1 pulldowns, its estimated ratio relative to the core proteins ranged from ∼1:3 to ∼1:10 depending on the measure used (see **Supplementary Information** and **Supplementary Table S5**). DARD1 includes a conserved rhodanese domain (IPR001763/PF00581), a member of the structurally related rhodanase-phosphatase superfamily (**Figure 2B**). Similar to DAPX1, DARD1 carries substitutions at conserved catalytic positions, arguing against canonical rhodanese activity, and outside the rhodanese region, the protein lacks recognizable counterparts in non-diplonemid organisms.

Substoichiometric partners of PPRD1 also included a class of proteins that we refer to as the DAHR (<u>D</u>eaminase <u>A</u>ssociated <u>H</u>elix–turn–helix <u>R</u>epeat) family, with at least nine members associated with the PPRD1–DAPX1 core (**Supplementary Table S5**). DAHR proteins lack recognizable catalytic domains and are characterized by conspicuous tandem arrays of ∼30–45-residue-long, antiparallel helix–turn–helix (rHTH) motifs; for example, DAHR1 (DIPPA_32813) contains 13 such motifs (**Figure 2B**). These motifs show no obvious sequence similarity to one another or to established α-helical repeat families, such as HEAT, tetratricopeptide repeat (TPR) or PPR. Like other α-solenoid proteins (see, e.g., [37]), DAHRs are predicted to adopt super-helical architectures. Structural similarity searches indicate that the *Diplonema* genome encodes at least 39 rHTH proteins, ∼85% of which are predicted to be mitochondrial [22]. Helix–turn–helix motifs of many α-solenoid proteins mediate contacts with proteins, RNA, or DNA [37,38]. DAHR proteins may therefore participate in RNA binding and/or protein–protein interactions, a possibility supported by the previous identification of two rHTH proteins (DIPPA_23652 and DIPPA_05352) as small mitoribosomal subunit assembly factors [35]. rHTH proteins are present in all examined diplonemids, but generally show low sequence conservation (see **Supplementary Information**).

The second notable multi-member group of substoichiometric partners was the DAPM (<u>D</u>eaminase <u>A</u>ssociated <u>P</u>PR <u>M</u>otif) family, characterized by PPR motifs but lacking conserved catalytic domains. The most abundant member, DAPM1 (DIPPA_70097; **Figure 2A,B**), contains seven tandem PPR motifs flanked by HTH motifs at its N and C termini (**Figure 2C**). For comparison, the inferred global proteome of *Diplonema* includes 121 high-confidence PPR proteins, 90% of which are predicted to be mitochondrial [22]. The detected protein domains and structural motifs of co-purified partner proteins are listed in **Supplementary Table S5B**. Taken together, the PPRD1–DAPX1 core associated with several classes of repeat-rich proteins with potential RNA– and/or protein-binding functions.

The combined LC-MS/MS data from the three baits were used to estimate protein–protein association scores (**Figure 2C; Supplementary Table S5**). A network integrating prey enrichment and prey-to-bait abundance ratios for all high-confidence partners revealed that PPRD1 and DAPX1 together formed the major hub of a 21-node network with 11 shared associated proteins (**Figure 2D**). Of the remaining significant partners, four associated specifically with PPRD1, one (an rHTH protein) with DAPX1, and two (both TPR proteins) with DAHR1. Most prey-to-bait abundance ratios varied among the three baits, except for the consistently robust PPRD1–DAPX1 association (**Supplementary Table S5A**). This pattern is compatible with a PPRD1–DAPX1 core that either transiently associates with accessory proteins or forms distinct co-existing complexes containing different sets of accessory proteins (see also **Supplementary Information**).

### 5. Structure prediction of the PPRD1–DAPX1 complex suggests possible roles for DAPX1

The predicted structure of the binary PPRD1–DAPX1 complex showed generally high local confidence scores for both subunits, whether modelled individually or as a complex (**Supplementary Figure S2,S6**). The predicted interface, supported by contacts conserved across diplonemid homologs (**Supplementary Figure S7**) and the regions of highest local confidence (**Supplementary Figure S8**), was dominated by helix–helix and helix–loop interactions (**Figure 2E**). All PPRD1 interface residues were located within the DYW domain, predominantly on the surface opposite the catalytic site (**Figure 2E,F**). The interacting residues of DAPX1 were more broadly distributed across the N-terminal three-quarters of the protein, primarily in the 8-kDa and palm subdomains of PolX (**Figure 2F**; **Supplementary Figure S7**). These residues were largely confined to the surface opposite the nucleic acid’s phosphate backbone-binding cleft, leaving the latter exposed. Thus, the DAPX1–PPRD1 interface could stabilize or regulate the deaminase domain, while the exposed PolX surface could provide an additional RNA-interaction site. These possibilities remain to be tested experimentally.

### 6. PPRD1 and DAPX1 silencing impairs RNA editing

To test whether PPRD1 and DAPX1 are involved in *Diplonema* mitochondrial RNA editing, we used gene silencing by RNA interference (RNAi) [40], applied here for the first time in *Diplonema*, whose nuclear genome encodes all required pathway components [41]. Readouts included cell proliferation, protein expression, and *in vivo* RNA editing. RNAi was induced by electroporating wild-type cells or tagged cell lines (PPRD1-PrA, DAPX1-PrA, or DAPX1-tdTomato; **Supplementary Table S3**) with 0.4–1.2 kbp-long double-stranded (ds) RNA molecules corresponding to the coding regions of target genes (see Methods, **Supplementary Tables S1,S2**).

We validated the RNAi approach by silencing the essential mitoribosomal protein gene MRPS63 (DIPPA_25715) [35]. Three days after dsRNA transfection, MRPS63-PrA levels dropped below 5% (**Figure 3A**). In wild-type cultures, silencing slowed cell division, followed by delayed proliferation. The culture was diluted (1:15) at day 3 post-transfection, and normal proliferation resumed 7–9-days later (**Figure 3B**). Similar growth inhibition and protein depletion were observed following the knockdown of either PPRD1, or DAPX1 (**Figure 3B–D**; **Supplementary Table S6**). No substantial cross-regulation of protein levels was detected: DAPX1 abundance changed only marginally upon PPRD1 silencing, and *vice versa* (**Figure 3C,D**). Controls included cultures transfected with similarly-sized dsRNA that did not target any *Diplonema* gene but had comparable nucleotide and codon composition, as well as mock-treated cultures. Both controls showed only negligible changes in cell proliferation and protein expression (**Figure 3B–D**).

**Figure 3.**
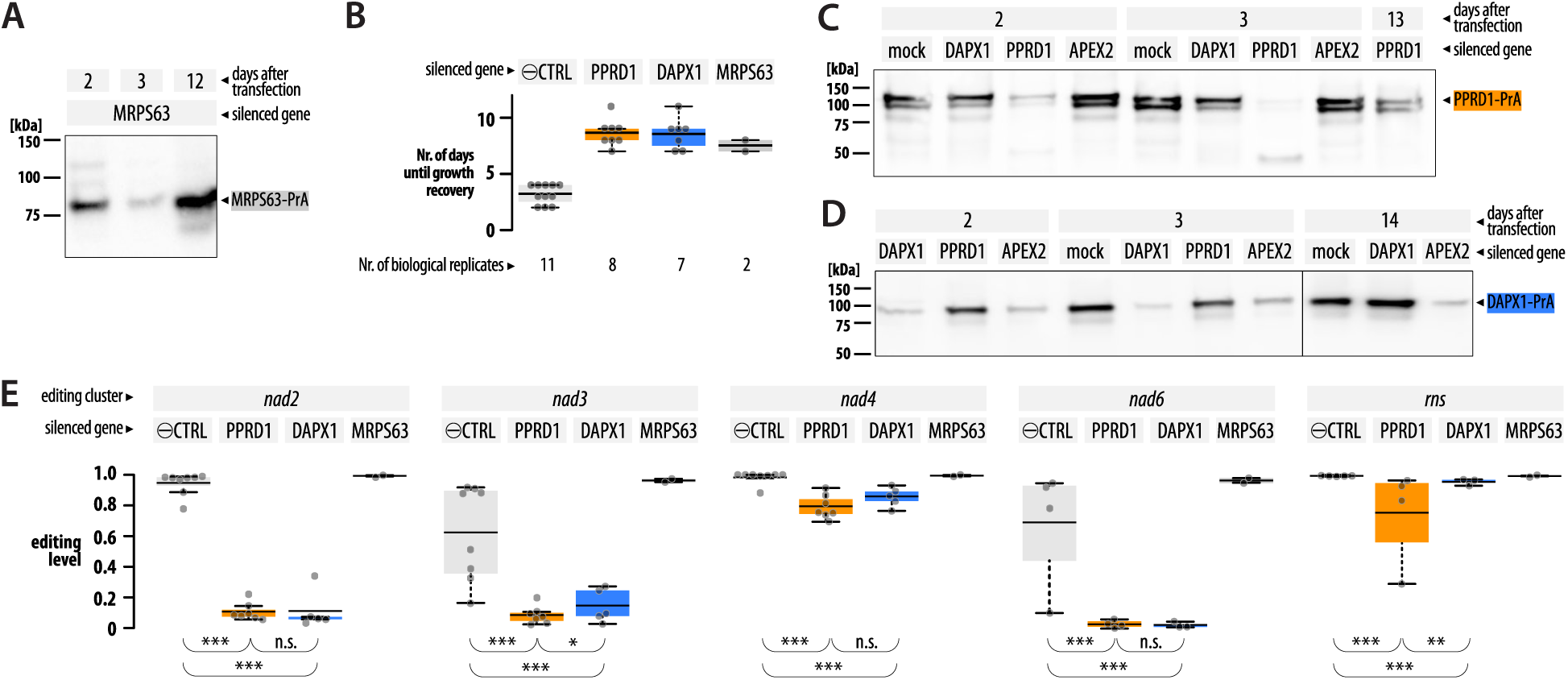
Transient RNAi-mediated silencing of PPRD1 and DAPX1 reduces cell proliferation, protein abundance, and mitochondrial RNA deamination. (**A**) Western blot analysis of PrA-tagged MRPS63 two, three, and 12 days after RNAi using dsRNA derived from the MRPS63 coding sequence. (**B**) Number of days until wild-type *Diplonema* cultures attained cell density typical of normal growth (following culture dilution three days after transfection). Samples: ⊖CTRL (combined negative controls including mock treatment, as well as dsRNAs targeting the PrA and APEX2 coding sequences, which are absent from wild-type *Diplonema*), PPRD1, DAPX1, and MRPS63 (dsRNAs targeting coding sequences of the corresponding genes). Box limits indicate the 25th and 75th percentiles, and centre lines show the mean. Each dot represents a biological replicate. (For details, see **Supplementary Table S6**.) (**C**, **D**) Western blot analysis of PrA-tagged PPRD1 (**C**) or DAPX1 (**D**) at various time points after silencing with dsRNAs derived from the coding sequences of DAPX1, PPRD1 or APEX2. The lower PPRD1-PrA band is a degradation product resulting from sample freezing and thawing. As the APEX2 dsRNA overlaps the tag region of the DAPX1-PrA fusion by 70 bp (but not that of PPRD1-PrA), APEX2 dsRNA treatment reduced DAPX1-PrA levels, but not PPRD1-PrA levels compared with the mock treatment. (**E**) RNA editing levels three days post-transfection, expressed as the proportion of fully edited reads among the sum of fully edited and pre-edited reads. Editing levels were determined separately for the five deamination clusters in *Diplonema* mitochondrial mRNAs: *nad2* (module m4), *nad3* (m2), *nad4* (m1), *nad6* (m1), *rns* (m1). Note that *nad2*, *nad3*, and *nad6* were initially referred to as *y3*, *y1*, and *y5*, respectively [39]. Samples, box limits, and centre lines are as in (**B**). Each dot represents a biological replicate. For statistical analyses, Fisher’s exact test was used for pairwise comparisons: n.s. (not significant), p > 0.01; *, 0.01 ≥ p > 10^−10^; **, 10^−10^ ≥ p > 10^−20^; ***, p ≤ 10^−20^. (For details, see **Supplementary Information**, **Supplementary Figure S9**, and **Supplementary Table S7**.)

To assess the effects of silencing PPRD1 or DAPX1 on *in vivo* deamination editing, we measured, by RT-PCR and nanopore sequencing, editing within clusters in five mitochondrial RNAs, four mRNAs encoding subunits of the NADH:ubiquinone dehydrogenase (*nad2*, *3*, *4*, *6*) and the small-subunit mitochondrial RNA (*rns*) collectively including a total of 27 A-to-I and 83 C-to-U sites [20,39]. (For more details about the experimental setup, see **Supplementary Information**, **Supplementary Figure S9**, **Supplementary Table S7**.) Silencing PPRD1 or DAPX1 increased the odds of observing fully pre-edited clusters (i.e., matching the genomic sequence) relative to fully edited clusters. The effect was most pronounced for *nad2* (∼200-fold) and smaller for the other transcripts (∼5–80-fold; **Figure 3E**; **Supplementary Table S7D,G**). Both knockdowns had similar effects on *nad2*, *nad4*, and *nad6*, whereas the deamination of *nad3* and *rns* was more affected by PPRD1 than DAPX1 silencing (**Supplementary Table S7F**). Importantly, the impact on RNA editing did not appear to result from growth inhibition *per se* because editing levels in MRPS63-silenced cultures remained unchanged (**Figure 3E**).

Together, these results show that i) RNA interference is gene-specific, with no evidence of substantial cross-regulation between PPRD1 and DAPX1; and ii) both PPRD1 and DAPX1 are required for deamination RNA editing in *Diplonema* mitochondria. Explaining the differences observed among transcripts and between PPRD1 and DAPX1 silencing will ultimately require establishing *in vitro* deamination assays with defined substrates.

## DISCUSSION

We showed here that *Diplonema* PPRD1, a mitochondrion-localized divergent homolog of PPR-DYW deaminases, associates with numerous mitochondrial proteins. Among these, DAPX1, a protein of unknown function carrying a non-catalytic DNA polymerase domain, co-purified with PPRD1 at an approximately 1:1 ratio. Down-regulation of either protein *in vivo* in *Diplonema* impaired mitochondrial deamination RNA editing of both C and A residues, indicating that the two partners are critical for the process.

A central question arising from our work is how a single PPR-DYW protein could drive highly specific RNA editing at numerous C-to-U and A-to-I sites. In plant systems, the multi-site problem is typically solved by multiple site-specific PPR proteins that carry their own DYW domain or recruit a catalytic DYW partner.

Likewise, all surveyed Discoba lineages other than diplonemids that retain PPR-DYW proteins contain multiple paralogs. In plant PPR-DYW proteins, the PPR tract provides much of the sequence specificity by binding RNA upstream of the edited nucleotide [29,30], although recent structural work shows that contacts involving the E1/E2 and DYW domains also contribute to positioning the target cytidine [11]. Diplonemid PPRD1 contains only seven P-type motifs, and the residue pairs at canonical RNA-recognition positions show limited correspondence to the plant PPR code. Although the PPR array of PPRD1 likely contributes to RNA binding, its residue pairs alone do not readily explain site specificity—let alone the recognition of more than 1000 editing sites in hemistasiid diplonemids [42]. Even if *Diplonema* PPRD1 targeted entire editing clusters rather than individual sites, it would still need to identify at least eight distinct positions within approximately 15 kb of the pre-edited transcriptome. However, previous analyses revealed no common sequence or structural features in the cluster-surrounding RNA that could serve as targeting signals [20]. These observations therefore favour a model in which PPRD1-associated proteins contribute substantially to site selection, as we discuss below. Site-resolved RNA-binding assays and targeted perturbation of candidate partners should help distinguish between these possibilities.

A second outstanding question is whether PPRD1 itself catalyzes RNA deamination. Its DYW domain contains residues characteristic of catalytically active DYW domains. Plant DYW domains are established C-to-U deaminases, as supported by structural studies and as demonstrated *in vitro* by editing with purified recombinant PPR-DYW proteins [6,7,11] and *in vivo* in heterologous systems [8,9]. Bacterial heterologous expression of chimeric proteins composed of unrelated PPR array and DYW domain has proven especially powerful for investigating the molecular mechanisms of deamination editing [10,13,32]. Although this latter approach has had a fair success rate with plant DYW proteins, swapping the DYW domain of *Diplonema* PPRD1 into moss editing factors yielded no detectable deamination (see also **Supplementary Information**). Ongoing activity assays with affinity-purified or recombinant PPRD1 preparations have likewise yet to show activity, and may require systematic optimization of assay conditions, including substrate design, protein combinations, cofactors, and conditions that preserve or reconstitute the native editing complex.

We could, nevertheless, demonstrate that gene silencing of PPRD1 reduces not only C-to-U, but also A-to-I editing in mitochondrial transcripts *in vivo* across all five examined mitochondrial RNA-editing clusters encompassing 110 deamination sites. Engineered APOBEC, ADAR, and TadA enzymes with broadened substrate specificity, but modest performance, have been developed [43–45]. However, to our knowledge, no naturally occurring enzyme with dual C– and A-deaminase activity has been described. PPRD1 thus appears to have an unusually broad catalytic capability.

Our study also revealed that PPRD1 does not operate on its own. The protein-protein interaction data point to a multi-component deamination machinery that parallels, but is not equivalent to, plant editosomes. In seed plants, it is not uncommon that RNA recognition and C-to-U catalysis are distributed among site-recognition PPR proteins and trans-acting DYW proteins [32,46–48], recently confirmed by reconstitution experiments [13]. Also widespread in seed plants are several families of novel non-PPR regulatory and assembly factors, such as MORF/RIP and ORRM, first identified in *Arabidopsis* [34,49,50]. Some act at multiple editing sites, whereas others affect editing more broadly. As a result, seed plant editing complexes vary substantially in composition across editing sites, not only in the PPR recognition factor but also in their associated catalytic and regulatory non-PPR components (e.g., [12,51,52]).

In *Diplonema*, DAPX1 co-purified with PPRD1 in near-equal proportions, and post-RNAi editing impairment supported a shared, functionally important PPRD1–DAPX1 core. Structural modelling suggested that DAPX1 may stabilize PPRD1 while leaving its PolX-like nucleic-acid-binding face and other surface regions exposed for potential RNA and protein interactions. The two proteins also associated with at least a dozen substoichiometric proteins with predicted RNA-binding properties, including members of multiple helical-repeat families. The DAHR and DAPM proteins are plausible candidates for contributing to the recognition of individual sites or editing-site clusters, and could help explain how a system dependent on a single PPR-DYW protein achieves the observed high target specificity. Our data are compatible with two possibilities: transient PPRD1–DAPX1 assemblies that exchange accessory factors, or distinct PPRD1–DAPX1 assemblies that contain different accessory factors. Together, our findings support a multipartite deamination-editing machinery centred on a PPRD1–DAPX1 core and provide a framework for dissecting the organization, target specificity, and evolutionary origins and diversification of this unusual RNA editing system in diplonemids.

## ACKNOWLEDGEMENTS

We thank Corinna Benz (University of South Bohemia, Czechia) for advice in designing RNA interference experiments, Dhruba Dey (Université de Montréal, Canada and Indian Institute of Science, India) for preliminary examinations of interactions in predicted protein structures, and Juliane Lenz (Heidelberg University, Germany) for preparing several PPRD1-chimeras used in the heterologous bacterial *in vivo* RNA deamination assays. We also thank Gerardo Ferbeyre and Daniel Zenklusen for sharing material (both at Université de Montréal, Canada), and Juan D. Alfonzo and Jesse Leavitt (Brown University, Providence, USA) for ongoing assistance with the development of *in vitro* RNA editing assays.

## AUTHOR CONTRIBUTIONS

G.B, M.V., conceptualization;

L.C.A., P.M.B., N.K., M.V., investigation;

G.B., N.K., B.F.L., M.V., formal analysis;

M.O., J.P., R.S., M.V., methodology;

G.B., project administration, supervision;

J.A., G.B., M.O., J.P., R.S., funding acquisition;

G.B., M.V., writing-original draft;

L.C.A., P.M.B., G.B., N.K., B.F.L., M.O., J.P., R.S., M.V., writing – review & editing.

All authors read and approved the final manuscript.

## DATA AVAILABILITY

New data reported in this article are available in the on-line **Supplementary Information**. New mass spectrometry proteomics data were deposited to the ProteomeXchange Consortium via the PRIDE partner repository [53] with the dataset identifier PXD080080. Updated *nad2*, *nad3*, *nad4* transcript sequences used in this study were deposited in GenBank under accession numbers KU341375, KU341373, and KU341367, respectively. Additional **Supplementary Material** (e.g., all scripts used for editing quantification and statistical analyses, *D. papillatum* proteomics-related resources, etc.) are archived at a FigShare repository (DOI: 10.6084/m9.figshare.33438424).

## AI USE DISCLOSURE

LLMs were used for language editing. In addition, generative AI tools were used to assist in parts of the statistical analysis workflow. All models, parameters, and outputs were reviewed, validated, and interpreted by the authors, who take full responsibility for the results.

## MATERIALS AND METHODS

For more details on experimental procedures, see the **Supplementary Information**.

### Cell cultivation

*Diplonema papillatum* (ATCC 50162) was cultivated axenically in saline liquid medium supplemented with horse serum as described earlier [39,41].

### DNA and RNA extraction, (RT-)PCR, and generation of cell lines

DNA was isolated using phenol-chloroform [54], and RNA was extracted with a Trizol substitute [55]. To reverse transcribe RNA and amplify DNA, we used Protoscript II and Q5 DNA polymerase (New England Biolabs), respectively. To generate *D. papillatum* cell lines expressing proteins C-terminally tagged with either Protein A (PrA), or a fluorescent protein, we inserted ∼1.0–1.6 kbp-long regions including part of the gene body and the 3′ UTR of the targeted gene as upstream and downstream homology arms, respectively, into plasmids carrying PrA, YFP or tdTomato tags and the neomycin resistance marker [55]. For heterologous expression in *Escherichia coli*, the open reading frames of PPRD1 and DAPX1 (lacking their predicted mitochondrial targeting pre-sequences) were inserted into pETDuet-1-based plasmids. Primers and plasmids are listed in **Supplementary Table S1,S2**. All *Diplonema*-tagging constructs were transformed by electroporation as described earlier and selected in a G418-containing medium [55,56]. Cell lines used in this study are listed in **Supplementary Table S3**.

### Fluorescence microscopy

The expression of tdTomato-tagged proteins in live cells and immunofluorescence-based localization of PrA-tagged proteins in formaldehyde-fixed cells were observed under an Eclipse Ts2R-FL microscope (Nikon), following recommendations for diplonemid imaging [57], with minor modifications.

### Isolation of mitochondria

Sub-cellular fractions, including mitochondria, were isolated by separating homogenized cells on a velocity sucrose gradient essentially as described [54]. Mitochondrial enrichment was gauged by the proportion of mitochondrial rRNAs and mtDNA separated by agarose gel electrophoresis. Detailed protocols are also available at https://www.protocols.io/researchers/matus-valach.

### Western blotting

Briefly, 10 µg of total proteins per lane of whole-cell lysate or gradient fraction samples were separated on 10% Tris-Tricine SDS-PAGE gels [58], electro-transferred onto a PVDF membrane, which was then blocked with 5% non-fat milk and incubated with a primary rabbit anti-PrA antibody (Sigma P3775, RRID: AB_261038; used at 1:10,000). Following the incubation with secondary goat anti-rabbit Horse Radish Peroxidase (HRP)-conjugated antibody (Cell Signaling Technology 7074, RRID: AB_2099233; used at 1:2,000), bands were visualized by chemiluminescence using a CCD-camera instrument.

### Protein A-mediated pulldown of native complexes and mass spectrometry

Single-step affinity purification (AP) and MS/MS was performed in 3–4 replicates as previously described [35], with minor modifications as detailed in Supplementary Information. Briefly, *Diplonema* whole-cell pellets were frozen in liquid nitrogen and pulverized with a cryo-mill, dissolved in the presence of salts and detergents (buffers detailed in **Supplementary Information**), and then incubated with rabbit IgG-conjugated magnetic beads (rabbit-IgG, Sigma I5006, RRID: AB_1163659; Dynabeads M-270 Epoxy). Captured proteins were on-bead-digested with trypsin, cleaned up with C18 ZipTip pipette tips (Millipore) and loaded on a C18 column installed in the Easy-nLC 1200 system (Proxeon Biosystems). The HPLC system was coupled to an Orbitrap Fusion mass spectrometer (Thermo Scientific) through a Nanospray Flex Ion Source with data acquisition using Xcalibur v4.0 and Tune v2.0. Datasets are described in **Supplementary Table S4**.

### Analyses of mass spectrometry and interactome data

Peptide-spectrum matching, protein identification, scoring, and spectrum counting, as well as ion-based quantifications were performed in Fragpipe v23.1 (https://github.com/Nesvilab/FragPipe) using MSFragger v4.3 [59], Philosopher v5.1.2 [60], and IonQuant v1.11.11 [61], respectively. To score protein–protein interactions, we used SAINTexpress [62] and fold-change calculations as implemented at the CRAPome database and analysis webserver [63] and visualized the results using the ProHits-viz tool suite [64]. Protein abundances across experiments were compared using spectral counts and precursor intensities after their normalization to the number of theoretically observable peptides by calculating PAI (Protein Abundance Index) and iBAQ (intensity-Based Absolute Quantification) values (**Supplementary Table S5**). Only proteins with SAINT probability score (SP) >0.9 and with average and geometric fold-change (FC_A and FC_B) >4 were considered as interaction partners of a particular bait. No abundance threshold was imposed to avoid discarding any proteins that interacted specifically, but weakly or at low substoichiometric ratios. The interactome graph was built using Cytoscape v3.10.4 [65].

### Gene silencing

Double-stranded RNA (dsRNA) for *Diplonema* gene silencing was produced by PCR-amplifying target regions (**Supplementary Tables S1, S2**) and cloning phosphorylated amplicons into pBluescript in both orientations. Selected plasmids were linearized with EcoRI and transcribed using T7 RNA polymerase. Following ssRNAs extraction, the two strands were annealed. Ten to 20 µg of purified dsRNA was used per transfection of late exponential-phase *Diplonema*. About 10^8^ cells were electroporated in the presence of dsRNA or, alternatively, water for mock controls. Treated cultures were allowed to recover before sampling two and three days after the transfection (RNA for RT-PCR, proteins for Western blotting, and cells for microscopy). Cultures were then further cultivated until normal growth resumed (**Supplementary Table S6**) and sampled again to gauge the silenced gene expression by Western blotting or microscopy.

### RNA-editing site profiling after gene silencing and statistical analysis

Total RNA was extracted from mock-treated or dsRNA-silenced (PrA, APEX2, MRPS63, PPRD1, DAPX1) wild type cells three days after transfection. Mitochondrial transcript regions undergoing deamination RNA editing (namely, the deamination clusters of *nad2*, *nad3*, *nad4*, *nad6*, and *rns* transcripts) were reverse transcribed and amplified by RT-PCR using gene-specific primers (**Supplementary Table S1**). Amplicons from the same sample were then combined and sequenced using the Oxford Nanopore technology at Plasmidsaurus. All experiments were performed in at least 3 biological replicates. Nanopore reads were mapped on the edited transcript reference sequences using Minimap2 v2.24 [66] and parsed with bedtools v2.31.1 [67] to include only amplicon-spanning reads. Editing rates were inferred from read counts by classifying reads as fully pre-edited or edited based on cluster-specific sequence patterns (see **Supplementary Information** and also **Supplementary Table S7A–E; Supplementary Figure S9**). Per-sample counts were summarized in contingency tables for control and silenced conditions and used to calculate odds ratios; confidence intervals and p-values were obtained using Fisher’s exact test (**Supplementary Table S7F**). Pairwise gene and cluster comparisons were performed using Wald z-tests on logistic regression coefficients, and cluster-specific silencing effects were evaluated with binomial logistic regression and likelihood-ratio tests (**Supplementary Table S7G–I**).

### Basic sequence analyses

Basic sequence analyses (including local sequence similarity search, domain identification, multiple sequence alignment, structural alignment and search, phylogeny reconstruction) followed the procedures established previously [68]. The following tools were used: HMMER [69], BLAST [70], Muscle5 [71], Clustal Omega v1.2.3 [72], MAFFT v7.490 [73], PROMALS3D [74], Foldseek v6 [75], PSIPRED (https://github.com/psipred/psipred), AIUPred [76], Deeploc2 v2.1 [77], WebLogo3 [78], FastTree v2.1.11 [79], IQ-TREE 2 [80]. Databases used: InterPro r107.0 [81], Pfam r38.0 [82], Uniprot [83], NCBI CDD [84], PPR – Pentatricopeptide Repeat Proteins [85] (https://ppr.plantenergy.uwa.edu.au), PDBe [86], *Diplonema* predicted foldome [87].

### Protein structure modeling and comparative analyses

To generate structural models of proteins and their complexes, we used AlphaFold v2 and v3 [88,89], after supplementing, as proposed previously [90], their default databases with eukaryotic sequences under-represented or missing from public databases. These included the EukProt collection [91] and proteomes inferred from transcriptomes of 12 diplonemid species [42]. Structures were visualized in UCSF ChimeraX v1.11 [92]. Three-dimensional structural comparisons and alignments were done with the ‘matchmaker ‘tool of ChimeraX and

FoldMason v4 [93]. Protein–protein interaction interfaces in the AlphaFold3-predicted complexes were determined using the PICKLUSTER plug-in of ChimeraX [94]. ZincSight [31] was used to predict and evaluate zinc ion binding sites.

## Abbreviations

ADAR: adenosine deaminase acting on RNA
AP: affinity purification
CDAT: cytidine deaminase acting on tRNA
LC–MS/MS: Liquid chromatography–tandem mass spectrometry
IP: immuno-precipitation
MTP: mitochondrial targeting pre-sequence
mtDNA: mitochondrial DNA
pHMM: profile Hidden-Markov model
pLDDT: predicted Local Distance Difference Test
PPR: pentatricopeptide repeat
PrA: Protein A
rHTH: repeated helix–turn–helix
RNAi: RNA interference
RNP: ribonucleoprotein
TPR: tetratricopeptide repeat

