## Supplemental Information for "A microeukaryotic PPR-DYW protein supports multisite C- and A-deamination RNA editing"

### SUPPLEMENTARY RESULTS AND DISCUSSION

#### Structural features of PPRD1

*PPR repeats*

Seminal work in *Arabidopsis* has subdivided PPR motifs into three classes, the P (standard, length 35 residues), L (long, length 36–37), S (short, length 30–33), and SS (very short, length 28–30). Plant PPR proteins that confer sequence-specific target recognition in organelle RNA editing are composed of a succession of P-L-S motifs [1], whereas some sequence-specific RNA-binding designer proteins consist solely of P motifs [2]. As in most non-plant lineages [3] (but see, e.g., [4]), PPR motifs in diplonemids belong to the P type [5]. The PPR array in *D. papillatum* PPRD1 protein includes seven putative motifs that share only 10–26% similarity among each other. Five of these motifs are 35-residue long with significant, yet low sequence similarity to plant P-type motifs. With 31 residues, the two central motifs fall in the length range of plant S motifs, but profile Hidden-Markov model (HMM) searches returned slightly better scores with P than S motifs. Across the diplonemids examined, the protein sequence of the PPR array is well conserved but conspicuously less in the central motifs that are in some cases shorter or longer by one residue compared those from *D. papillatum* (**Supplementary Figure S1**). Taken together, the two central, short PPR motifs exhibit accelerated rates of sequence change compared to the five flanking P motifs. The biological significance is yet unclear.

According to the PPR code established in plant proteins, an array of seven PPR motifs can theoretically recognize a specific RNA septamer target, as two residues within each PPR-motif, position 5 and 35, collectively determine base specificity. For example, the residue pairs T-N, (T/A)-D, and D-N recognize A, G, and C, respectively, in RNA [6,7]. In contrast, in the PPR array of *Diplonema* PPRD1, the residues at positions 5 and 35 only match these canonical base-recognition pairs to a limited degree and in all cases (N-D, N-N, N-L) are known to display preferential affinity for pyrimidines, resulting in the inference of a degenerate motif 5′–?YYr?Yy–3′ (upper case, high preference; lower case, low preference; ?, no known preference). Similar results were obtained with all twelve diplonemids we examined (see **Supplementary Figure S1**). There is also a possibility that the PPR array of *Diplonema* PPRD1 does not mediate sequence-specific RNA recognition at all. In this regard, it could resemble PPR proteins found in mitochondrial ribosomes, which generally lack residues for sequence discrimination. Ribosomal PPR proteins typically interact with RNA either primarily via the phosphate backbone, or engage their repeats in protein–protein interactions (reviewed in [8]). While PPRD1 likely binds RNA, the precise functional role of its PPR array has yet to be established.

*DYW domain*

The C-terminal region of PPRD1 harbours an approximately 120 residue-long DYW domain (PF14432/IPR032867), which belongs to the superfamily of cytidine deaminase-like domains. It includes all the function-critical residues that bind two $Zn^{2+}$ ions: the canonical catalytic $Zn^{2+}$ (shared by all cytidine deaminase folds), as well as the second $Zn^{2+}$ unique to DYW deaminases, which appears to stabilize the domain [9]. The domain was named after its fairly conserved Asp-Tyr-Trp (DYW) tripeptide motif [10]. This tripeptide is TGW in *D. papillatum* and DGW or EGW in the other examined diplonemids. Notably, the Tyr, characteristic of plant DYW domains and corresponding to residue 959 of the Arabidopsis reference protein OTP86 (PP296_ARATH) is replaced by a Gly in all examined diplonemids. Another conspicuous difference to canonical plant DYW domains concerns the second zinc-binding motif. In all studied diplonemids, the first invariable Cys typical of plant DYW proteins (position 954 in the reference), is substituted by an Asp, and an additional residue (leucine, lysine, or methionine) is inserted immediately after this Asp (see **Supplementary Figure S1**). Whether this substitution and the adjacent inserted residue may confer PPRD1's expanded substrate repertoire remains to be determined.

The AlphaFold3-predicted three-dimensional (3D) structure of diplonemid PPRD1 has high local confidence scores (pLDDT >80) across the major portion (~70%) of the protein. Low-confidence segments are confined to the very N-terminus, predicted disordered loops, a 10 residue-long insertion connecting the beta sheet and α1 helix of the deaminase domain, and the 14 residue-long diplonemid-specific C-terminal extension (**Supplementary Figure S2**). The bulk of the modelled *Diplonema* DYW domain aligns well with the crystal structure of the *Arabidopsis* DYW (RMSD=0.834 Å across homologous residues, excluding the 'gating' sub-domain and the C-terminal extension; **Figure 1C, Supplementary Figure S3**). The main structural divergence lies in a region referred to in *Arabidopsis* proteins as the 'gating' sub-domain (or 'DYW deaminase insertion'), which undergoes conformational changes proposed to regulate the active site [9,11]. Such flexible, dynamic regions are inherently difficult for AlphaFold to model accurately [12,13].

**PPR-DYW proteins show lineage-specific retention and expansion across Discoba**

We examined PPR-DYW copy number and diversification across Discoba. Malawimonads were included as a phylogenetically informative deep-branching and slowly evolving counterpart [14] and plant homologs served as outgroups. Complete or nearly complete inferred proteomes are available for the malawimonads *Malawimonas jakobiformis*, *M. californiana*, and *Gefionella okellyi* (B.F. Lang, unpublished), and the discobids *Andalucia godoyi* (Jakobida; [15]), *Acrasis kona* (Heterolobosea; [16]), *Euglena gracilis* (Euglenida; [17]), and *Trypanosoma brucei* (Kinetoplastida; [18]). Using a profile hidden Markov model built from aligned diplonemid PPRD1 sequences, we searched these proteomes for PPR-DYW homologs. No homologs were detected in *M. jakobiformis*, *Andalucia*, *Euglena*, or *Trypanosoma*, whereas multiple paralogs were identified in *M. californiana*, *Gefionella*, and *Acrasis*. In contrast, diplonemids consistently encode a single PPR-DYW homolog.

A phylogenetic tree of the retrieved proteins, together with diplonemid PPRD1 sequences and representative plant homologs as the outgroup, broadly followed the expected species relationships [14] (**Supplementary Figure S4**). Within individual lineages, multiple homologs formed distinct paralog groups, consistent with lineage-specific expansions of the PPR-DYW family. The distribution among malawimonads is consistent with ancestral presence followed by secondary loss in the *M. jakobiformis* lineage and expansion in *M. californiana* and *Gefionella*. Similarly, the multiple *Acrasis* homologs indicate expansion in heteroloboseans, whereas the absence of detectable homologs from the examined jakobid, euglenid, and kinetoplastid is consistent with lineage-specific loss. Thus, the single-copy state of diplonemid PPRD1 homologs strongly contrasts with the expansions observed in these lineages and in plants (see [1]).

**Proteins co-purified with PPRD1, DAPX1, and DAHR1**

*Subcellular localization*

Out of the 21 proteins that were significantly enriched across the immuno-affinity pulldowns with each of the three baits (PPRD1-PrA, DAPX1-PrA, DAHR1-PrA), 17 were computationally predicted to be mitochondrial with high confidence [5], with the remaining four proteins classified as low-probability candidates. More specifically, DIPPA_20138 (interacting with all three baits) and DIPPA_00956 (partner exclusively of DAHR1) were classified as possibly mitochondrial, though with low scores. DIPPA_24282 and DIPPA_30118, both specific partners of PPRD1, had higher probability of being cytosolic than mitochondrial. However, counterparts of the former in other diplonemids were systematically predicted to localize to the mitochondrion (with relatively high scores); hence we regard DIPPA_24282 as mitochondrial, too.

*rHTH proteins*

Proteomes of all examined diplonemids include a number of repeated helix–turn-helix (rHTH) proteins, classified as such based on their predicted structural features, i.e., predominantly composed of series of anti-parallel α-helices, each ~14–22 amino acid residues-long, yet with little to no significant sequence similarity in their structural repeats. The DAHR class represents a subset of rHTH proteins associated with the deaminase core (represented by PPRD1 and DAPX1). Sequence conservation across diplonemid species is generally low, as is taxonomic distribution. For example, DAHR1 (DIPPA_32813) has a recognizable homolog only in *Lacrimia lanifica* and *Sulcionema speckii*, and DIPPA_31654 in *D. japonicum*, *D. aggregatum*, and *D. ambulator*; whereas DIPPA_22109 and DIPPA_26365 have no clear counterparts. With 30.6% pairwise identity, the only diplonemid-wide and highly conserved DAHR protein is DIPPA_24282. However, its abundance, enrichment, and interaction scores were only significant in PPRD1-PrA pulldowns (**Supplemental Table S5**).

*TPR proteins*

Interpro searches classified the two most significant proteins in the DAHR1 pulldowns, DIPPA_00956 and DIPPA_33632, as tetratricopeptide repeat (TPR) proteins. However, this assignment was purely sequence-based as the repeated structural motifs are 42 residues long, rather than 34 expected for TPR motifs. DIPPA_00956 had clearly recognizable homologs only in four out of 11 species (*D. japonicum*, *D. aggregatum*, and *D. ambulator*, and *L. lanifica*), while DIPPA_33632 was fairly conserved across all diplonemids. The latter protein has a distant paralog in *D. papillatum* (TR118232__c0_g1_i1__m.27569) and other diplonemids, but also a similarly distant homolog in dinoflagellates. This suggests a conserved, yet unknown function. As dinoflagellate mitochondria and plastids carry out RNA editing of various kinds [19–21], the presence of a factor common to these two distant protist lineages is not implausible. It remains to be determined if DIPPA_33632 plays a conserved role in mitochondrial gene expression.

*Protein partners with previously assigned functions*

DIPPA_21168, an rHTH-class member, was the only significant protein in the pulldowns (with PPRD1-PrA) with previously assigned function: it represents the mitoribosomal protein MRPS51 (mS51) of the mitochondrial small subunit (mtSSU) [22]. As MRPS51 is an abundant protein, the detected interaction with PPRD1 could be spurious. Alternatively, it may have dual functions, acting both as a structural mitoribosomal protein and as an editing factor. Notably, in *Trypanosoma*, mS51 interacts to a limited extent with mtSSU rRNA [23]. It is conceivable that DIPPA_21168 also interacts with mtSSU rRNA, which, with 45 sites in a 374 nt-long molecule, represents the most extensively deamination-edited transcript in *Diplonema* [22,24]. It would be worthwhile to test whether DIPPA_21168 is the specificity factor for editing pre-mtSSU rRNA or is implicated, through its RNA-binding capacity, in linking deamination editing and mitoribosome formation.

**Strategies to assess the RNA deamination activity of PPRD1**

To study otherwise refractive DYW domains of plant RNA deaminases, chimeric PPR-DYW strategy has proved particularly productive [11,25,26]. We therefore attempted to test the PPRD1 deaminase activity by swapping its DYW-domain into the mitochondrial editing factors PPR56 and PPR65 from the moss *Physcomitrium patens*. Specifically, the *Diplonema* domain was fused at various, structurally compatible positions to a region downstream of PPR arrays of these editing factors, and the activity was assayed using a heterologous *in vivo* editing test in bacteria by RT-PCR, as originally devised in [25], with or without the co-expression of DAPX1. Using the same experimental setup, we also modified—by mutating various amino acid residues—the moss DYW domain to make it resemble that of *Diplonema*. However, in neither case did we detect the expected C-to-U conversions (nor A-to-I conversions), whereas the native plant proteins (used as positive controls) efficiently edited the programmed site at a 50–100% rate in our setup. Evidence from more recent studies focusing on the effects of chimeric plant PPRD-DYW proteins showed that the long-range interplay between the catalytic domain and the RNA binding region is generally complex, with apparent non-trivial, variable (positive or negative) effects of DYW domains on the RNA recognition and/or binding by the PPR arrays [27,28]. Thus, diplonemid DYW domain might be evolutionarily too distant from the plant proteins to support effective cooperation between the chimera moieties.

### SUPPLEMENTARY MATERIALS AND METHODS

#### Cell cultivation

*Diplonema papillatum* (ATCC 50162) was cultivated axenically without shaking at 15–22 °C in ocean salt medium (OS) containing 33 g/L Instant Ocean Sea Salt (Instant Ocean) and supplemented with 1–2% (v/v) horse serum as described earlier [29,30], and with chloramphenicol (40 mg/L) to prevent bacterial contamination. For mitochondrial isolation, yeast extract was added to the medium to 0.04% (w/v). For the selection and preservation of the tagged cell lines, the antibiotic G418 was added at 100–200 mg/L.

#### DNA and RNA extraction, (RT-)PCR, plasmid construction, and generation of *Diplonema* cell lines

DNA was isolated using phenol-chloroform [31], and RNA was extracted with a Trizol substitute [32]. To reverse transcribe (RT) RNA and amplify DNA, we used Protoscript II and Q5 DNA polymerase (New England Biolabs), respectively. We generated *D. papillatum* cell lines expressing proteins that are C-terminally tagged with either Protein A (PrA), or the fluorophore tdTomato. To this end, we inserted using NEBuilder approach (New England Biolabs) ~1.0–1.6 kbp-long regions that included part of the gene body (upstream homology arm) or the 3′ UTR (downstream arm) of the targeted gene into the plasmids pDP006 (PrA tag), pDP015 (PrA) or pDP036 (tdTomato) that carry tags and a neomycin resistance marker (see [32,33] and this study). For heterologous expression in *Escherichia coli*, open reading frames of PPRD1 and DAPX1 (lacking their predicted mitochondrial targeting pre-sequences) were inserted into pETDuet-1-based plasmids. All cloning procedures were done in chemically competent [34] *E. coli* DH5α cells. Primers and plasmids are listed in **Supplementary Table S1, S2**.

*Diplonema* gene-tagging constructs were transformed by electroporation as described earlier and transformants were selected in G418-containing medium [32,35,36]. Expression of the tagged proteins was confirmed by Western blotting. Generation of the cell line expressing mtPrA was described previously [35]. Cell lines used in this study are listed in **Supplementary Table S3**.

#### Fluorescence microscopy

Fluorescent samples were inspected using an Eclipse Ts2R-FL microscope (Nikon) following, with minor modifications, the recommendations for diplonemid imaging [37]. To determine the subcellular localization of PrA-tagged proteins by immunofluorescence, we harvested 15 mL culture of cells from late exponential phase at 2,000 ×g, washed the cells once in fresh OS medium, then fixed them in 4% formaldehyde (in OS), for 20 min at 20 °C. Following a spin at 4,000 ×g to remove the crosslinker, cells were permeabilized in pre-chilled methanol for 30 min at −20 °C. Following another spin at 4,000 ×g, fixed and permeabilized cells were washed twice in 1×PBS, then blocked in 1×PBS supplemented with 0.1% Tween-20 and 3% BSA by incubation on the rotator at 20 °C for 1h. The blocking buffer was removed by centrifugation and primary anti-PrA antibody (Sigma P3775, RRID: AB_261038; 1:1,000–1:2,000) was added in fresh blocking buffer. After a 14h-long incubation on the rotator at 4 °C, cells were centrifuged and washed 2–3 times in 1×PBS. Samples were then incubated with goat anti-rabbit Alexa Fluor 568-conjugated secondary antibody (Invitrogen A-11036, RRID: AB_10563566; 1:500) in the blocking buffer, on the rotator in the dark for 2h at 20 °C. Labelled cells were washed 3 times in 1×PBS, then resuspended in 1×PBS supplemented with DAPI (1:2,000 final dilution, ThermoFisher Scientific), transferred on a slide, covered, and then imaged. Photos were processed in NIS-Elements AR v5.02 (Nikon) and overlayed in Affinity Photo v2.6.5 (Serif Europe).

#### Western blotting

After determining the protein concentration in each sample using the Bradford dye-binding method, whole-cell lysates or gradient fractions were separated by Tris-Tricine SDS-PAGE [38]. Ten µg of total protein dissolved in the loading buffer were dispensed per lane (unless otherwise stated, i.e., **Supplementary Figure S5**). After electrophoresis on a 10% polyacrylamide gel, separated proteins were electro-blotted onto a 0.45-µm PVDF membrane (Amersham Hybond, Cytiva) in a Tris-Glycine pH9.2-transfer buffer using a Mini Trans-Blot cell (Biorad). Following a one hourlong blocking step at 20 °C in 5% fat-free milk in 1× TBS (supplemented with Tween-20 to 0.1%), blots were incubated in 1× TBST with the primary anti-PrA antibody (Sigma P3775, RRID: AB_261038, used at 1:10,000) for 16–20 h at 4 °C. After several washes in 1× TBST, blots were incubated with secondary goat anti-rabbit Horse Radish Peroxidase (HRP)-conjugated antibody (Cell Signaling Technology 7074, RRID: AB_2099233; used at 1:2,000) for 1 h at 20 °C. Bands were visualized by chemiluminescence using Clarity Max Western ECL substrate (Bio-Rad) or SignalFire ECL Plus reagent (Cell Signaling Technology) on a ChemiDoc MP instrument (Bio-Rad).

**Protein A-mediated pulldowns**

Single-step affinity purification (AP) was performed in multiple replicates (four for PPRD1 and three each for DAPX1 and DAHR1) as described previously [22]. *Diplonema* cell pellets from cultures harvested in late exponential phase were frozen in liquid nitrogen and ground by cryo-milling. Hundred-milligram aliquots of cell powder were vortexed in 500 µL of the AP buffer (for PPRD1 and DAPX1: 30 mM Tris-HCl pH7.6, 20 mM KCl, 25 mM $MgCl_2$; for DAHR1: 30 mM Tris-HCl pH7.6, 20 mM KCl) supplemented with protease inhibitor cocktail (Mini-complete EDTA-free, Roche) and 1 % detergent (PPRD1 and DAPX1: Triton X-100; DAHR1: dodecylmaltoside). After a brief sonication (20 W, 2 sec) and lysate clearing by centrifugation (16,100×g, 4 °C, 10 min), the cleared lysate was topped up to a volume of 900 µL with the detergent-free AP buffer to adjust the detergent's final concentration to 0.1%. Relative abundance of each PrA-tagged bait in the cleared lysates was determined by Western blot and then used to determine the volume required for affinity purifications so as to saturate 3.75 mg of Dynabeads M-270 Epoxy conjugated in advance with rabbit IgG (Sigma I5006, RRID: AB_1163659). Magnetic beads prewashed in the AP buffer with 0.1% detergent were combined with the cleared lysates for a 30-min incubation on a rotator at 4 °C. After six washes in the AP buffer with 0.1% detergent, the beads were washed once in the AP buffer with 0.01% detergent for 5 min with agitation and then four times in 30 mM Tris-HCl pH7.6 and 5 mM $MgCl_2$. This was followed by on-bead tryptic digestion.

**Mass spectrometry**

Mass spectrometry was performed essentially as described previously [22]. Overnight on-bead digestion of affinity-purified (AP) complexes was performed using 500 ng trypsin in 20 mM Tris-HCl pH8.0 at 37 °C. The digestion was stopped by adding formic acid to a final concentration of 2%. Tryptic peptides were speedvac-dried and stored at −80 °C. Prior to LC-MS/MS, protein digests were re-solubilized under agitation for 15 min in 10 µL of 0.2% formic acid. Desalting/cleanup of the digests was performed by using C18 ZipTip pipette tips (Millipore). Eluates were dried down in a vacuum centrifuge and then re-solubilized under agitation for 15 min in 10 µL of 1% ACN and 1% formic acid. Samples were loaded into a 75 µm i.d. × 150 mm Self-Pack C18 column installed in the Easy-nLC 1200 system (Proxeon Biosystems).

The samples were then processed at the Institut de recherches cliniques de Montréal (IRCM) mass spectrometry facility as follows. The buffers used for chromatography were 0.2% formic acid (buffer A) and 100% acetonitrile with 0.2% formic acid (buffer B). Peptides were eluted with a two-slope gradient at a flowrate of 250 nL/min. Solvent B was first increased from 1 to 34% during 74 min and then from 34 to 93% during another 8 min. The HPLC system was coupled to Orbitrap Fusion mass spectrometer (Thermo Scientific) through a Nanospray Flex Ion Source. Nanospray and S-lens voltages were set to 1.4 kV and 60 V, respectively. Capillary temperature was set to 250 °C. Full scan MS survey spectra (m/z 360-1550) in profile mode were acquired in the Orbitrap with a resolution of 120,000 and a target value of 4e5. The 20 most intense peptide ions were fragmented in the HCD collision cell and analyzed in the linear ion trap with a target value at 1e4 and a normalized collision energy at 30 V. MS3 scanning was performed upon detection of a neutral loss of phosphoric acid (48.99, 32.66 or 24.5 Th) in MS2 scans. Target ions selected for fragmentation were dynamically excluded for 25 sec after two MS2 events. For data acquisition, we used Xcalibur v4.0 and Tune v2.0. Proteomics datasets reported here are described in **Supplementary Table S4**.

**Analyses of mass spectrometry and interactome data**

Spectrometry data in Thermo RAW format were converted to mzML using ThermoRawFileParser v1.4.5 [39]. Searches for peptide-spectrum matches, protein identification, scoring, and spectrum counting, as well as ion-based quantifications were conducted from within Fragpipe v23.1 (https://github.com/Nesvilab/FragPipe) using MSFragger v4.3 [40], Philosopher v5.1.2 [41], and IonQuant v1.11.11 [42], respectively. The Fragpipe workflow configuration, sample manifest with assignments of PrA-tagged bait (PPRD1, DAPX1, DAHR1) and control (mtPrA) replicates, and the proteome file are provided in **Supplementary Material** (see DOI: 10.6084/m9.figshare.33438424 and 10.6084/m9.figshare.28635440). To score protein-protein interactions (PPIs), we used SAINTexpress [43] implemented at the CRAPome database and analysis webserver [44] and visualized the results using the ProHits-viz tool suite [45]. Only proteins with SAINT probability score (SP) >0.9 and with both average and geometric fold-change scores (FC_A and FC_B) >4 were considered interaction partners of a particular bait. To avoid discarding any proteins interacting specifically but weakly or at low sub-stoichiometric ratios, we did not impose any abundance threshold. Protein abundance across experiments was compared using spectral counts and precursor intensities after their normalization to the number of theoretically observable peptides by calculating PAI (Protein Abundance Index) and iBAQ (intensity-Based Absolute Quantification) values (**Supplementary Table S5A**). Observable peptides were determined as previously reported [29] using the MS-Digest tool of ProteinProspector v6.8.1 (https://prospector.ucsf.edu/prospector/mshome.htm). Briefly, based on the prevalence and distribution of peptides and their ions in the proteomics datasets (**Supplementary Material**; DOI: 10.6084/m9.figshare.33438424), for each protein, we considered as theoretically observable all peptides that were within the m/z range of 350–1,500, with +2 and +3 charges, and no variable modifications; these parameters covered >95% of all identified peptide-spectrum matches. The normalized abundances and SAINTexpress fold changes and scores were then combined for the purpose of visualization and categorization of identified PPI candidates. The interactome graph was generated using Cytoscape v3.10.4 [46].

**Gene silencing by RNA interference**

Double-stranded RNA (dsRNA) was used to trigger gene silencing in *Diplonema* cells. To generate dsRNA, we PCR-amplified with gene-specific primers (**Supplementary Table S1**) selected regions of various targets (PrA, APEX2, MRPS63, PPRD1, DAPX1; **Supplementary Table S2**) using as template plasmids or RT reactions of RNA extracted from the corresponding *Diplonema* cell lines. Phosphorylated amplicons were then inserted into pBluescript vector (digested by EcoRV and dephosphorylated by Quick-CIP) using Blunt-TA master mix (all enzymes New England Biolabs) and introduced into *Escherichia coli* DH5α cells. Colonies were screened by extracting plasmids using Monarch spin miniprep kit (New England Biolabs) followed by appropriate restriction enzyme digestions to determine insert orientation. For each target *Diplonema* gene, plasmids with a forward and a reverse insert were then linearized by EcoRI digestion, phenol:chloroform-extracted, and transcribed separately using the HiScribe T7 High Yield RNA Synthesis Kit (New England Biolabs). The produced complementary transcripts were extracted using the Trizol substitute (see above) and the amount of ssRNA was assessed by denaturing gel electrophoresis and spectrophotometry. Complementary ssRNAs (50 µg in forward and reverse orientation each) were annealed in 50 mM Tris pH7.0, 200 mM NaCl after denaturation at 85 °C and allowed to slowly cool to 50 °C; the dsRNA was precipitated and again analyzed by gel electrophoresis and spectrophotometry to verify quality and quantity. Fifteen to 20 µg of dsRNA were then used per *Diplonema* transfection depending on the size of the transcript.

Prior to electroporation, *Diplonema* cells from a culture grown in OS with 1% horse serum in late exponential phase were harvested by centrifugation (2,000 ×g, 5 min), then washed once in the OS medium and once the RNAi buffer (90 mM $NaH_2PO_4$, 6 mM KCl, 0.15 mM $CaCl_2$, 50 mM HEPES, pH 7.3). Per transfection, ~$10^8$ cells were resuspended in the RNAi buffer supplemented with dsRNA or water as a control, and exposed to an electric pulse (1,500V, 0.8 ms). Fresh medium was added immediately afterwards, and the suspension was transferred to a culture flask to allow cell recovery. RNA for RT-PCR, protein for Western blotting, and cells for microscopy were sampled 2 and 3 days after transfection (2/3 and 14/15 volume of the culture, respectively), with the culture topped up each time to the initial volume with fresh medium. Cultures were then left to recover under standard cultivation conditions until cells resumed normal growth as determined through bright-field microscopy observations and only then sampled again; this usually took 2–21 days depending on the target gene and cell line (**Supplementary Table S6**). Cell cultures were passaged at least once more to confirm regular growth pattern. If a culture did not recover within 42 days after the transfection, it was discarded.

**RNA-editing site profiling after gene silencing**

Three days post-transfection, total RNA was extracted from wild type cells that were mock-treated or gene-silenced. At this timepoint, Western blotting of PrA-tagged cell lines showed that the target protein in the culture had decreased to <5% of the level under normal growth conditions, while cell morphology and behaviour were not visibly affected. Note that in wild-type cells, the dsRNAs containing parts of PrA and APEX2 coding sequences lacked any target. Thus, in total, eight cultures served as negative controls: three were water-treated (*0neg*), three were transfected with PrA dsRNA (*A-v1*) and two with APEX2 dsRNA (*A-v2*). The remaining cultures were transfected with dsRNA targeting genes for silencing: seven cultures with PPRD1 dsRNA (*B-v1*, *B-v2*, *B-v3*) and five with DAPX1 dsRNA (*C-v1*) (see **Supplementary Table S2**).

To investigate the impact of gene-silencing on deamination RNA editing, we reverse transcribed the RNA with an anchored oligo-dT (or oligo-dA) primer and then PCR-amplified regions spanning the editing clusters of *nad2*, *nad3*, *nad4*, *nad6*, and *rns* transcripts using gene-specific primers and the anchored oligonucleotide (**Supplementary Table S1**; **Supplementary Figure S9**). (Note that originally, *nad2*, *nad3*, and *nad6* were provisionally referred to as '*y3*', '*y1*', and '*y5*' [29].) To increase the amplification specificity of *rns*, which is the only single-exon mitochondrial transcript of *Diplonema* and thus especially prone to contamination by immature (pre-edited) precursors and by mtDNA, we leveraged the fact that the mature *rns* has a 3′ U-tail and a free hydroxyl at its 3′ terminus [24]. The RNA sample was therefore A-tailed using *E. coli* poly-A polymerase (New England Biolabs) and then reverse transcribed using the CDSIII oligonucleotide as any regular mRNA (see **Supplementary Table S1**). Only those amplicons that contained both the natural U tail and the artificially added A-tail were taken into account in the editing site profiling. Similar to other *Diplonema* mature mitochondrial mRNAs, *nad6* carries a poly-A tail; however, it also contains a 50 U-long stretch between two of its exons [24], interfering with reliable RT-PCR across the region. To increase the *nad6* RT-PCR efficiency, instead of the poly-A tail-targeting oligonucleotide CDSIII, we used dp137, which is complementary the U-tract downstream of the transcript's editing cluster (**Supplementary Table S1**; **Supplementary Figure S9**).

Amplicons were separated on agarose gels, extracted, quantified, and those from the same culture pooled at equimolar ratios (~1 ng/µL per 100 bp). Samples were then submitted for the Standard PCR Premium sequencing at Plasmidsaurus (USA) using the Oxford Nanopore technology, which generated ~500–1,500 reads per amplicon. All experiments were performed in at least three biological replicates.

Nanopore reads from each sample were mapped on the edited amplicon reference sequences using Minimap2 v2.24 [47] with a k-mer size of 12, which improved mapping of pre-edited reads compared to the default setting: `minimap -x map-ont -k 12 --frag=yes --secondary=no -a refSeq.fasta input -o output.sam`. Using samtools v.1.22.1, SAM files were converted to BAM, while removing unmapped reads: `samtools view -b -F 4 output.sam > output.bam`. Mapped reads were next filtered with bedtools v2.31.1 [48] using default parameters, which selected those that overlapped by at least 1 nt both the upstream (5′) RT-PCR primer and the last 10 nt at the 3′ end of the RT-PCR amplicon: `bedtools intersect -r -abam input.bam -b position_dn.bed > out_dn.bam ; bedtools intersect -r -abam out.bam -b position_up.bed > out_dn_up.bam`.

The resulting BAM alignments were then used to infer RNA editing rates as follows. Each sample was assigned to one of two conditions: 'control' or 'silenced'. Editing rates were inferred from read counts (**Supplementary Table**

**S7A–E**). Reads were classified as pre-edited or edited by exact matching to the following cluster-specific diagnostic sequence patterns:

- editing cluster of *nad2*, pre-edited: `CTGACGTGGTATGCTGTCCC`
- editing cluster of *nad2*, edited: `TTGGTGTGGTATGTTGTTTT`
- editing cluster of *nad3*, pre-edited: `CACGTGGGTGGATGCAGTCATCCCTAC`
- editing cluster of *nad3*, edited: `TGTGTGGGTGGGTGTGGTTGTTTCTAT`
- editing cluster of *nad4*, pre-edited: `CACAGCCTGTGCTGGATGATCCCACCATACCTAACAGTACATCCACCTCGGCCTC`
- editing cluster of *nad4*, edited: `TGTGGTTTGTGTTGGATGATTTTATTGTATTTGGTAGTGTATTTGTTTTGGTTTT`
- editing cluster of *nad6*, pre-edited:
  `CTACCTGTAGCCACGTGGTGTACTGCTGGTATCAGCGTGCATGTGCCATCTGCTGTACCATGCGTACCTGC`
- editing cluster of *nad6*, edited:
  `TTATTTGTAGTTATGTGGTGTACTGCTGGTATCAGTGTGTATGTGTTATTTGTTGTATTATGTGTATTTGT`
- editing cluster of *rns*, pre-edited:
  `CCCAGCTGGGTATGTGCTCTGTACCGCTGTACGGTACTAGCTATCCATCCACGCAGCTACGTCCACCATGCTGTGCTATCAGCCC`
- editing cluster of *rns*, edited:
  `TTTGGTTGGGTGTGTGTTTTGTGTTGTTGTGTGGTGTTGGTTGTTTGTTTGTGTGGTTGTGTTTGTTGTGTTGTGTTGTTGGTTT`

Counts were obtained from aligned reads by parsing BAM records and counting exact matches to patterns (shown here for one pattern file): `samtools view out_dn_up.bam | grep -F -c -f patternfile` . Reads excluded by this step typically contained partially edited sequences and/or sequencing errors within the diagnostic cluster.

Per-sample counts of fully pre-edited and fully edited reads were compiled into contingency tables by condition ('control' vs. 'silenced') and separately for each transcript (*nad2*, *nad3*, *nad4*, *nad6*, *rns*) using the custom script `bam_to_contingency.pl` (**Supplementary Table S7A–E**). One PPRD1-silenced biological replicate (V2VFW7_5_WT-0805-76h-B-v2) showed an unusually high proportion of pre-edited *rns* reads relative to the other replicates. However, excluding the outlier sample had no material effect on the pairwise odds ratio comparisons (see below), indicating that the observed gene-specific differences were not driven by this sample.

**Statistical analysis of RNA editing rates after gene silencing**

In brief, the statistical analysis included odds ratio calculations via pooled Fisher 2×2 tests and logistic regression for testing interactions using the R environment.

*Odds ratios and confidence intervals for each silenced gene and examined transcript*

The question addressed here was: for each gene and transcript, did gene silencing increase the odds that a read was pre-edited (versus edited)? The effect size was expressed, for each 'gene × transcript' combination, as a silenced/control odds ratio, computed directly from pooled 2×2 edited vs pre-edited read-count tables. For counts pooled across samples, see **Supplementary Table S7A–E**, bottom part.

The odds ratio (OR) was defined as: `OR = (P_s / E_s) / (P_c / E_c)` , where P_s, E_s are the counts of pre-edited and edited reads in 'silenced' and P_c, E_c the corresponding counts in 'control' conditions. The OR and support values were calculated with a custom script using as input the above generated contingency tables. Transcript-dependent silencing effects were evaluated among transcripts analyzed from the same set of biological samples, notably *nad2*, *nad3*, and *nad4*, using the command: `Rscript compare_genes_across_transcripts.R contingency-nad2.tsv contingency-nad3.tsv contingency-nad4.tsv`. (Silencing effects were not compared for *nad6* and *rns*, because their sets of biological replicates differed.) The first output file of this script, `OR_by_gene_transcript.tsv`, reported for each 'gene × transcript' combination the odds ratio, 95% confidence interval (CI_low / CI_high), and p_value, the latter two from Fisher's exact test. In addition, the output file printed four summary metrics for coherence checking and interpretation: pre-edited fraction in controls: `(P_c / (P_c+E_c))`; pre-edited fraction in silenced: `(P_s / (P_s+E_s))`; and total control reads: `P_c+E_c`, as well as total silenced reads: `P_s+E_s` (**Supplementary Table S7F**).

*Testing of transcript-specific effects of gene silencing*
The question addressed by this test was whether the effect of silencing a given gene impacted RNA editing of the transcripts differently. For that, we tested whether the silenced/control odds ratios were equal across transcripts using a likelihood-ratio test. For each gene, a binomial logistic regression model was fitted to per-sample counts, where the log odds of a read being pre-edited were modeled as a function of condition ('control' vs 'silenced'), transcript identity, and sample (controlling for sample-specific baseline differences). A second model included in addition a 'condition × transcript' interaction term. A likelihood-ratio test comparing these two models assessed whether the silencing effect differed between transcripts. The interaction p-value, reporting evidence that the effect varied depending on a particular transcript, was calculated with the script `compare_genes_across_transcripts.R` as above. The second output file created by the script, `interaction_test_by_gene.tsv` (**Supplementary Table S7G**), described the transcript dependence, and contained two columns: 'gene' and 'interaction_p'.

*Pairwise comparisons of effects on edited transcripts and silenced genes*
Two types of comparisons were performed. The first addressed the question: For a given gene knockdown, do two edited transcripts respond differently? For this estimation, we calculated the log(OR) and evaluated the differences using Wald z-tests using the following command: `Rscript pairwise_transcript_effects.R PPRD1 contingency-nad2.tsv contingency-nad3.tsv [etc.] > PPRD1_pairwise.tsv` and `Rscript pairwise_transcript_effects.R DAPX1 contingency-nad2.tsv contingency-nad3.tsv [etc.] > DAPX1_pairwise.tsv`. The results from two executions for either PPRD1 or DAPX1 were compiled in **Supplementary Table S7H**.

The second question addressed was: For a given edited transcript, does knockdown of PPRD1 have a different effect compared to that of DAPX1? For this calculation, we used, e.g., `Rscript pairwise_or_differences.R contingency-nad2.tsv > pairwise_OR_differences-nad2.tsv`. The results from the executions with inputs `contingency-nad2.tsv`, `contingency-nad3.tsv`, etc., are compiled in **Supplementary Table S7I**.

**Basic protein sequence analyses**
Multiple protein-sequence alignments (MSAs) were performed with the HMM-based Muscle5 [49], Clustal Omega v1.2.3 [50], MAFFT v7.490 [51], and PROMALS3D [52]. Conserved protein domains were searched in InterPro release 107.0 [53] (https://www.ebi.ac.uk/interpro) and NCBI CDD [54]. Profile HMM searches using Pfam 38.0 models [55] or plant PPR-motif subclasses P, P1, P2, L1, L2, S1, S2, SS [56] (https://ppr.plantenergy.uwa.edu.au) were performed with the HMMER3 suite [57]. Secondary structures and disordered regions were predicted using PSIPRED (https://github.com/psipred/psipred) and AIUPred [58]. Searches for structural similarities were performed using Foldseek v6 [59], using the in-house (see below) and previously published [60] databases of predicted diplonemid structures, as well as experimental structures from PDBe [61]. Searches for proteins with repeated helix–turn–helix (rHTH) structural motifs were done using predicted structures of the 9 identified DAHR proteins as queries; matches with scores >100 and E-values <0.0025 were classified as rHTH-like proteins. Sub-cellular localization of *D. papillatum* proteins was determined previously [5]; for additional diplonemid proteins, we used Deeploc2.1 [62].

**Phylogenetic analysis**
Using a profile HMM built from the diplonemid DYW domain, we searched for homologous domain sequences in proteome or translated transcriptome datasets from other species obtained through the NCBI Genome or NCBI Nucleotide databases. Species and accession numbers are as follows: *Physcomitrium patens* (NCBI Genome: GCF_000002425.5), *Euglena gracilis* strain Z (GCA_039621445.1; Eugr-Chen2024.verA1.pep.fa), *Naegleria gruberi* (GCF_000004985.1), *Naegleria lovaniensis* (GCF_003324165.1), *Heterolobosea* sp. BB2 (GEZU00000000.1; transcriptome, translated in-house), *Pharyngomonas kirbyi* (GECH00000000; transcriptome, translated in-house), *Acrasis kona* (GCA_026419775.2), and *Trypanosoma brucei brucei* TREU927 (GCF_000002445.2). The complete

proteomes of *Andalucia godoyi*, *Malawimonas jakobiformis*, *Malawimonas californiana*, and *Gefionella okellyi* (B.F. Lang & S. Prince, unpublished), which were also used in this analysis are provided in **Supplementary Material** (DOI: 10.6084/m9.figshare.33438424).

The shared DYW domain was aligned with MUSCLE5 [49]. Positions not aligned by hmmalign with posterior of probability PP = 1.0 [57] were removed, leaving 172 amino acid positions. Phylogenetic analysis was performed with IQ-TREE 2 and automatic model selection (`iqtree2 -s infile.phy -m MFP -B 1000 -alrt 1000 -nt AUTO`) [63]. The resulting tree was displayed in FigTree (https://github.com/rambaut/figtree).

**Protein structure modeling and comparative analyses**

For structure modeling, we used AlphaFold2 [12] and AlphaFold3 [13]. As performance is influenced by the depth of a query protein's MSA, particularly for proteins from evolutionarily distant organisms, we supplemented AlphaFold's default databases with a custom dataset of sequences following a previously proposed approach [64]. Protein sequences from >1,000 eukaryotes under-represented or missing from public databases were added by combining sequences from various euglenozoans [64], the EukProt collection [65], and 12 diplonemid species [66] into a single FASTA file, clustered using MMseqs2 [67] at a 90% sequence identity threshold, then catenated to the default AlphaFold databases. Because our analysis required modelling interactions of a protein with multiple partners, using AlphaFold directly would have resulted in redundant searches against the sequence databases. To address this, we employed AlphaPullDown v1.0.4 [68], which performed the database search for each protein only once. For model predictions, we used default parameters and pre-trained model weights, without external structural restraints. Five candidate structures were generated and internally ranked by AlphaFold based on confidence metrics (e.g., mean predicted Local Distance Difference Test [pLDDT] and predicted aligned error [PAE]); only the top-ranked model for each protein was selected for downstream analyses. Structures were visualized and further examined in UCSF ChimeraX v1.11 [69] and PyMOL (https://www.pymol.org). Additional PDB file manipulations were done with pdb-tools v2.4.8 [70].

Multiple protein sequence-structure alignments were performed using the PROMALS3D webserver (http://prodata.swmed.edu/promals3d/promals3d.php) [52] and the results visualized in Jalview v2.11.5.1 [71]. To compare three-dimensional structures of proteins across species, we used the 'matchmaker' tool of ChimeraX, FoldMason v4 [72], and US-align v20241108 [73], all with default parameters. Protein-protein interaction interfaces in the AlphaFold3-predicted complexes were determined using the PICKLUSTER plug-in of ChimeraX [74] at a cut-off distance of 4.0 Å for grouping into spatially distinct clusters while excluding from interface calculations residues with pLDDT score <50. Only residues predicted to be in contact in at least ten out of 12 diplonemid species were considered for downstream analyses as conserved interaction sites. To predict zinc ion binding sites in the catalytic domain of PPRD1 (alone or in the predicted complex with DAPX1), we employed ZincSight [75] via the interactive Google Colab notebook (https://colab.research.google.com/github/MECHTI1/ZincSight/blob/master/ZincSight.ipynb); default parameters and settings were maintained and only the top-scoring site was considered for downstream analyses.

All custom code was executed using Python v3.11.8. More specifically, Bio.PDB, Bio.SeqIO, and Bio.AlignIO modules from the Biopython library were used to parse mmCIF/PDB files, write and edit structure files, and handle sequence and alignment data. The PyMOL Python API was used to programmatically generate and manipulate PyMOL environments. Pandas (https://pandas.pydata.org) and Matplotlib (https://matplotlib.org) were used for data handling and analysis, as well as to generate visualizations and graphs, respectively.

### SUPPLEMENTARY TABLES

**Supplementary Table S1**. Oligonucleotides used in this study.

| Oligo ID | Sequence (5′ → 3′) | Target(s) [a] | Application |
|---|---|---|---|
| CDS-III | ATTCTAGAGGCCGAGGCGGCCGACATGT[29]VN | *polyA* | RT-PCR |
| CDS-III-ter | ATTCTAGAGGCCGAGGCG | *nad2, nad3, nad6, rns* | RT-PCR |
| dp137 | ATTCTAGAGGCCGAGGCGGCCGACATGA[12] | *nad6* | RT-PCR |
| dp243 | CATCATACTGCTACCTACGACGGT | *nad4* | RT-PCR |
| dp264 | CTCAGGAGCACCTGGGCGTA | *nad3* | RT-PCR |
| dp268 | GGGTATGTGATGTCCTGGTGGTG | *nad4* | RT-PCR |
| dp274 | CTATGCAGTAGCCTGAGTAGCACG | *nad2* | RT-PCR |
| dp585 | TATGCTGTGGTACATGTGGTGG | *rns* | RT-PCR |
| dp615 | CGTAGTAGGTGGAGCTGTGTA | *nad6* | RT-PCR |
| dp373 | GCTTCCCATATGTGCGTCAGC | *APEX2* tag | PCR, dsRNA |
| dp375 | GGTGGGAACAACGAGGCAC | *pBA3295, pDP015* | PCR, tagging |
| dp384 | AAGAGAAGCTTGCCAGTAGTTGTTGAGAGT | *PPRD1* | PCR, tagging |
| dp385 | AGAGGTAACCTTTTGCTCAGTCCG | *PPRD1* | PCR, tagging |
| dp390 | GGTGAGCAAGACATCGCCAT | *PPRD1* | PCR, tagging & dsRNA |
| dp392 | AGTAGGAATTCTGGCTGACACCATTTGCG | *PPRD1* | PCR, tagging |
| dp393 | AATATGGATCCGAGTCCACCGCCATACTG | *PPRD1* | PCR, tagging |
| dp394 | AACGACTCCCAGGCACCC | *PrA tag* | PCR, dsRNA |
| dp395 | AAGTTTCTGGGCCTCTCCG | *pDP002, pDP006, pDP015* | PCR, tagging |
| dp403 | TGGGAAACGTGGCTGGAAG | *DAPX1* | PCR, dsRNA |
| dp418 | ATGCTGCAGTACATCGCCG | *PPRD1* | PCR, overexpression |
| dp419 | GAGTCCACCGCCATACTGAC | *PPRD1* | PCR, dsRNA |
| dp420 | ATGGCGATGTCTTGCTCACC | *PPRD1* | PCR, dsRNA |
| dp429 | CGGACCCGCTTCCCATAT | *PrA tag* | PCR, dsRNA |
| dp448 | AGTGCTGGACATCGGGGAATATTGAAGAGAGGGCCCAAACTCG | *DAPX1* | PCR, tagging |
| dp449 | AACCTGCGCTTCCAGGGGATCCCTGCGGGTCGTCGTCATT | *DAPX1* | PCR, tagging & dsRNA |
| dp450 | TTACGTGCTGCAAGTTTATACCAACGCCATGCTACC | *DAPX1* | PCR, tagging |
| dp451 | TAACAATTTCACCAAAGAATATTGATGAGACGGCCTTCC | *DAPX1* | PCR, tagging |
| dp452 | CCAACACGCTTCACGCCGAA | *DAPX1* | PCR, tagging |
| dp459 | GAAGTGCTGGACATCGCGGTCTGCAAGGAATGGTG | *DAHR1* | PCR, tagging |
| dp460 | AACCTGCGCTTCCAGGGGATCCAGTGAATCGAACGCCATCCC | *DAHR1* | PCR, tagging |
| dp461 | TTACGTGCTGCAAGTTTAGGGTGATTTTACGCTGGAAG | *DAHR1* | PCR, tagging |
| dp462 | TAACAATTTCACCAAAGAAGAAGTGGACGGGGAGAAGA | *DAHR1* | PCR, tagging |
| dp463 | CGGTCTGCAAGGAATGGTG | *DAHR1* | PCR, tagging |
| dp464 | CGAAGTGGACGGGGAGAAGA | *DAHR1* | PCR, tagging |
| dp520 | GAAGTGCTGGACATCGAATTCCGCTGCACTACATCCAT | *MRPS49* | PCR, tagging |
| dp521 | AACCTGCGCTTCCAGGGGATCCCATCGCGTCCCACTTCTGG | *MRPS49* | PCR, tagging |
| dp522 | TTACGTGCTGCAAGTTTGAAAAGTGCCCCTCTGTCG | *MRPS49* | PCR, tagging |
| dp523 | TAACAATTTCACCAAAGAATCGTGTCTGGAAGGATGAA | *MRPS49* | PCR, tagging |
| dp524 | GCTCCATTTGTTCCCGCCC | *MRPS49* | PCR, tagging |
| dp525 | GAAGTGCTGGACATCGAATTCGGTCTTCGCCGTGAGA | *MRPS63* | PCR, tagging |
| dp526 | AACCTGCGCTTCCAGGGGATCCCTCGCTACCGAGTTCGTCA | *MRPS63* | PCR, tagging |
| dp527 | TTACGTGCTGCAAGTTTACTGGACAAGAAGCAGAAGG | *MRPS63* | PCR, tagging |
| dp528 | TAACAATTTCACCAAAGAATTCATCACCGCCATCGTAG | *MRPS63* | PCR, tagging |
| dp529 | TAGACACCGGGGCATCATG | *MRPS63* | PCR, tagging & dsRNA |
| dp530 | ACCGTCGGCAATCCAGTTAT | *MRPS63* | PCR, tagging |
| dp557 | ATATAGGATCCAAGAGGAGATGCCGGGC | *PPRD1* | PCR, overexpression |
| dp558 | TACACAAGCTTAGAGTCCACCGCCATACTG | *PPRD1* | PCR, overexpression |
| dp559 | AGTATCATATGGCTTCGTCCTGCAACCA | *DAPX1* | PCR, overexpression |
| dp560 | ATATTCTCGAGTTACTGCGGGTCGTCGTCA | *DAPX1* | PCR, overexpression |
| dp562 | ATGGCTGCTGCCCATATGTATATCTCCTTCTTAAAGTTAAACAAA | *pETDuet, PPRD1* | PCR, overexpression |

| Oligo ID | Sequence (5′ → 3′) | Target(s) [a] | Application |
|---|---|---|---|
| dp579 | TTCGACGGGAAAAAGCTGAT | *PPRD1* | PCR, dsRNA |
| dp584 | ACATAGAATTCGGGGACTCCTTTGGACGA | *PPRD1* | PCR, overexpression |
| dp598 | ATGCATACCTCGGGCGACTACAAGG | *APEX2 tag* | PCR, dsRNA |
| dp599 | GGCGCCGGACGACGCC | *pDP015* | PCR, tagging & dsRNA |
| dp601 | GCAGTGAGCAAGGGCGAG | *pBA3295* | PCR, tagging |
| dp612 | TACCGCTCGCACCCTTCG | *pBA3295* | PCR, tagging |
| dp613 | CCTTAGTCAAGTGGATCTTGGTT | *pBA3295* | PCR, tagging |
| dp614 | TTTACATGAAGTCCACGCGGCC | *pDP015* | PCR, tagging |

[a] Base vectors used for the generation of tagging plasmids were previously reported in [22,32,33,35,36]. See also **Supplementary Table S2**.

**Supplementary Table S2**. Plasmids and other DNA constructs used in this study.

| **Plasmid ID** | **Target** | **Application** | **Derived from** |
|---|---|---|---|
| pBlu-RNAi-PPRD1v1-fwd | *PPRD1* | dsRNA production | pBluescriptII(SK+) |
| pBlu-RNAi-PPRD1v1-rev | *PPRD1* | dsRNA production | pBluescriptII(SK+) |
| pBlu-RNAi-PPRD1v2-fwd | *PPRD1* | dsRNA production | pBluescriptII(SK+) |
| pBlu-RNAi-PPRD1v2-rev | *PPRD1* | dsRNA production | pBluescriptII(SK+) |
| pBlu-RNAi-PPRD1v3-fwd | *PPRD1* | dsRNA production | pBluescriptII(SK+) |
| pBlu-RNAi-PPRD1v3-rev | *PPRD1* | dsRNA production | pBluescriptII(SK+) |
| pBlu-RNAi-DAPX1v1-fwd | *DAPX1* | dsRNA production | pBluescriptII(SK+) |
| pBlu-RNAi-DAPX1v1-rev | *DAPX1* | dsRNA production | pBluescriptII(SK+) |
| pBlu-RNAi-MRPS63v1-fwd | *MRPS63* | dsRNA production | pBluescriptII(SK+) |
| pBlu-RNAi-MRPS63v1-rev | *MRPS63* | dsRNA production | pBluescriptII(SK+) |
| pBlu-RNAi-APEX2v1-fwd | *APEX2 tag* | dsRNA production | pBluescriptII(SK+) |
| pBlu-RNAi-APEX2v1-rev | *APEX2 tag* | dsRNA production | pBluescriptII(SK+) |
| pBlu-RNAi-PROTAv1-fwd | *PrA tag* | dsRNA production | pBluescriptII(SK+) |
| pBlu-RNAi-PROTAv1-rev | *PrA tag* | dsRNA production | pBluescriptII(SK+) |
| pDP006 | n.a. | base vector for *Diplonema* tagging | pDP002 (PrA) |
| pDP015 | n.a. | base vector for *Diplonema* tagging | pDP006 (PrA) |
| pDP006B | *PPRD1* | *Diplonema* tagging | pDP006 (PrA) |
| pDP015C | *DAPX1* | *Diplonema* tagging | pDP015 (PrA) |
| pDP015E | *DAHR1* | *Diplonema* tagging | pDP015 (PrA) |
| pDP015S | *MRPS49* | *Diplonema* tagging | pDP015 (PrA) |
| pDP015T | *MRPS63* | *Diplonema* tagging | pDP015 (PrA) |

**Supplementary Table S3**. *Diplonema papillatum* cell lines used in this study.

| Cell line ID | Tagged gene | Application | C-terminal tag [c] |
|---|---|---|---|
| WT-072017 [a] | none | immuno-fluorescence microscopy, RNAi | not applicable |
| 10A-D30 [b] | *MPRA1* [b] | Western blot, proteomics | mt-PrA |
| 06B-B11 | *PPRD1 (DIPPA_21441)* | Western blot, proteomics, immuno-fluorescence microscopy, RNAi | 10aa-PrA |
| 06B-A22 | *PPRD1 (DIPPA_21441)* | Western blot, immuno-fluorescence microscopy, RNAi | 10aa-PrA |
| 15C-A75 | *DAPX1 (DIPPA_70088)* | Western blot, proteomics, RNAi | 40aa-FLAG-WELQut-PrA |
| 15C-E25 | *DAPX1 (DIPPA_70088)* | Western blot, proteomics | 40aa-FLAG-WELQut-PrA |
| 15C-E61 | *DAPX1 (DIPPA_70088)* | Western blot, immuno-fluorescence microscopy, RNAi | 40aa-FLAG-WELQut-PrA |
| 15E-E36 | *DAHR1 (DIPPA_32813)* | Western blot, proteomics | 40aa-FLAG-WELQut-PrA |
| 15E-E55 | *DAHR1 (DIPPA_32813)* | Western blot, proteomics | 40aa-FLAG-WELQut-PrA |
| 15S-F42 | *MRPS49 (DIPPA_31280)* | Western blot | 40aa-FLAG-WELQut-PrA |
| 15T-E32 | *MRPS63 (DIPPA_25715)* | Western blot, immuno-fluorescence microscopy, RNAi | 40aa-FLAG-WELQut-PrA |

[a] Wild-type *Diplonema papillatum* ATCC 50162, sub-clone used as the primary source for genome sequencing [30].
[b] A *D. papillatum* cell line carrying a genome-integrated construct that ectopically expresses Protein A fused to a mitochondrial targeting presequence [35].
[c] 10aa linker sequence is 'GSGSGSGSGS' ; 40aa linker sequence is 'GSPGSAGSGASGSGSASSGASAAGSSGASSGASAGSGA SA'.

**Supplementary Table S4**. Proteomics samples and datasets reported in this study.

| Sample ID | Source material | Detergent and $MgCl_2$ [b] | File ID |
|---|---|---|---|
| D76 [a] | mtPrA cells, replicate 1 | Triton X-100, 25 mM $MgCl_2$ | OF_20210531_OEF_06 |
| D81 [a] | mtPrA cells, replicate 2 | Triton X-100, 25 mM $MgCl_2$ | OF_20210716_OEF_01 |
| D82 [a] | mtPrA cells, replicate 3 | Triton X-100, 25 mM $MgCl_2$ | OF_20210716_OEF_02 |
| D83 [a] | mtPrA cells, replicate 4 | Triton X-100, 25 mM $MgCl_2$ | OF_20210716_OEF_03 |
| P27 | mtPrA cells, replicate 5 | Triton X-100, 25 mM $MgCl_2$ | OF_20230119_OEF_02 |
| P28 | mtPrA cells, replicate 6 | dodecylmaltoside, 25 mM $MgCl_2$ | OF_20230119_OEF_05 |
| P42 | mtPrA cells, replicate 7 | dodecylmaltoside, without $MgCl_2$ | OF_20250227_OEF_07 |
| P45 | mtPrA cells, replicate 10 | Triton X-100, 25 mM $MgCl_2$ | OF_20251202_OEF_03 |
| P46 | mtPrA cells, replicate 11 | Triton X-100, 25 mM $MgCl_2$ | OF_20251202_OEF_04 |
| P49 | mtPrA cells, replicate 12 | dodecylmaltoside, without $MgCl_2$ | OF_20251202_OEF_07 |
| P50 | mtPrA cells, replicate 13 | dodecylmaltoside, without $MgCl_2$ | OF_20251202_OEF_08 |
| D96 | PPRD1-PrA cells, replicate 4 | Triton X-100, 25 mM $MgCl_2$ | OF_20210531_OEF_02 |
| D97 | PPRD1-PrA cells, replicate 1 | Triton X-100, 25 mM $MgCl_2$ | OF_20210714_OEF_04 |
| D98 | PPRD1-PrA cells, replicate 2 | Triton X-100, 25 mM $MgCl_2$ | OF_20210714_OEF_05 |
| D99 | PPRD1-PrA cells, replicate 3 | Triton X-100, 25 mM $MgCl_2$ | OF_20210714_OEF_06 |
| P30 | DAPX1-PrA cells, replicate 1 | Triton X-100, 25 mM $MgCl_2$ | OF_20230119_OEF_01 |
| P43 | DAPX1-PrA cells, replicate 2 | Triton X-100, 25 mM $MgCl_2$ | OF_20251202_OEF_01_3 |
| P44 | DAPX1-PrA cells, replicate 3 | Triton X-100, 25 mM $MgCl_2$ | OF_20251202_OEF_02_2 |
| P41 | DAHR1-PrA cells, replicate 1 | dodecylmaltoside, without MgCl2 | OF_20250227_OEF_06 |
| P47 | DAHR1-PrA cells, replicate 2 | dodecylmaltoside, without MgCl2 | OF_20251202_OEF_05 |
| P48 | DAHR1-PrA cells, replicate 3 | dodecylmaltoside, without MgCl2 | OF_20251202_OEF_06 |

[a] Previously published data [22], PRIDE accession nr. PXD039927.
[b] Common buffer components: 30 mM Tris-HCl pH7.6, 20 mM KCl.

**Supplementary Table S5**. [**Excel file**] Interactome data of Protein A-tagged *Diplonema papillatum* PPRD1, DAPX1, and DAHR1. (**A**) Quantitative reports of bait-specific prey abundances, enrichments, and interaction scores of proteins identified in affinity purifications of PrA-tagged PPRD1, DAPX1, and DAHR1. Used to distinguish true interactions from non-specific associations. (**B**) Domain description, taxonomic distribution, and predicted sub-cellular localization of homologs for each significant component (SP > 0.9) in the affinity purification experiments.

**Supplementary Table S6**. Growth recovery of *Diplonema* wild-type cell cultures after transfection with gene-silencing dsRNA (**Figure 3B**).

| Date | dsRNA [a] | Suffix [b] | Recovery time [days] |
|---|---|---|---|
| 09 Sep 2025 | mock | 0neg | 6 |
| 09 Sep 2025 | PPRD1v1 | B-v1 | 11 |
| 09 Sep 2025 | PPRD1v2 | B-v2 | 11 |
| 09 Sep 2025 | DAPX1v1 | C-v1 | 11 |
| 17 Sep 2025 | mock | 0neg | 7 |
| 17 Sep 2025 | PPRD1v1 | B-v1 | 12 |
| 17 Sep 2025 | DAPX1v1 | C-v1 | 12 |
| 20 Sep 2025 | mock | 0neg | 5 |
| 20 Sep 2025 | PPRD1v2 | B-v2 | 12 |
| 20 Sep 2025 | DAPX1v1 | C-v1 | 12 |
| 27 Sep 2025 | mock | 0neg | 5 |
| 27 Sep 2025 | mtPrAv1 | A-v1 | 5 |
| 27 Sep 2025 | PPRD1v1 | B-v1 | 12 |
| 27 Sep 2025 | DAPX1v1 | C-v1 | 12 |
| 30 Sep 2025 | mock | 0neg | 6 |
| 30 Sep 2025 | APEX2v1 | A-v2 | 6 |
| 30 Sep 2025 | PPRD1v2 | B-v2 | 14 |
| 30 Sep 2025 | DAPX1v1 | C-v1 | 14 |
| 10 Nov 2025 | mock | 0neg | 7 |
| 10 Nov 2025 | APEX2v1 | A-v2 | 7 |
| 10 Nov 2025 | PPRD1v1 | B-v1 | 10 |
| 10 Nov 2025 | DAPX1v1 | C-v1 | 10 |
| 17 Nov 2025 | APEX2v1 | A-v2 | 7 |
| 17 Nov 2025 | PPRD1v3 | B-v3 | 11 |
| 21 Nov 2025 | MRPS63v1 | T-v1 | 10 |
| 24 Nov 2025 | mtPrAv1 | A-v1 | 7 |
| 24 Nov 2025 | DAPX1v1 | C-v1 | 10 |
| 24 Nov 2025 | MRPS63v1 | T-v1 | 11 |

[a] dsRNA was generated by annealing forward and reverse oriented transcript from pBluescript-derived templates (described in **Supplementary Table S2**).
[b] Sample suffix used to label RT-PCR sequencing runs, see **Supplementary Table S7**.

**Supplementary Table S7**. [**Excel file**] Effects of gene silencing on RNA editing. (**A–E**) Counts of fully edited or pre-edited reads comprising deamination RNA editing clusters in *nad2*, *nad3*, *nad4*, *nad6*, and *rns* mRNAs after gene silencing of PPRD1 and DAPX1. The counts served as the input data for all contingency-table and logistic-regression analyses. (**F–I**) Statistical analyses of gene silencing effects on the examined transcripts.

### Supplementary Figures

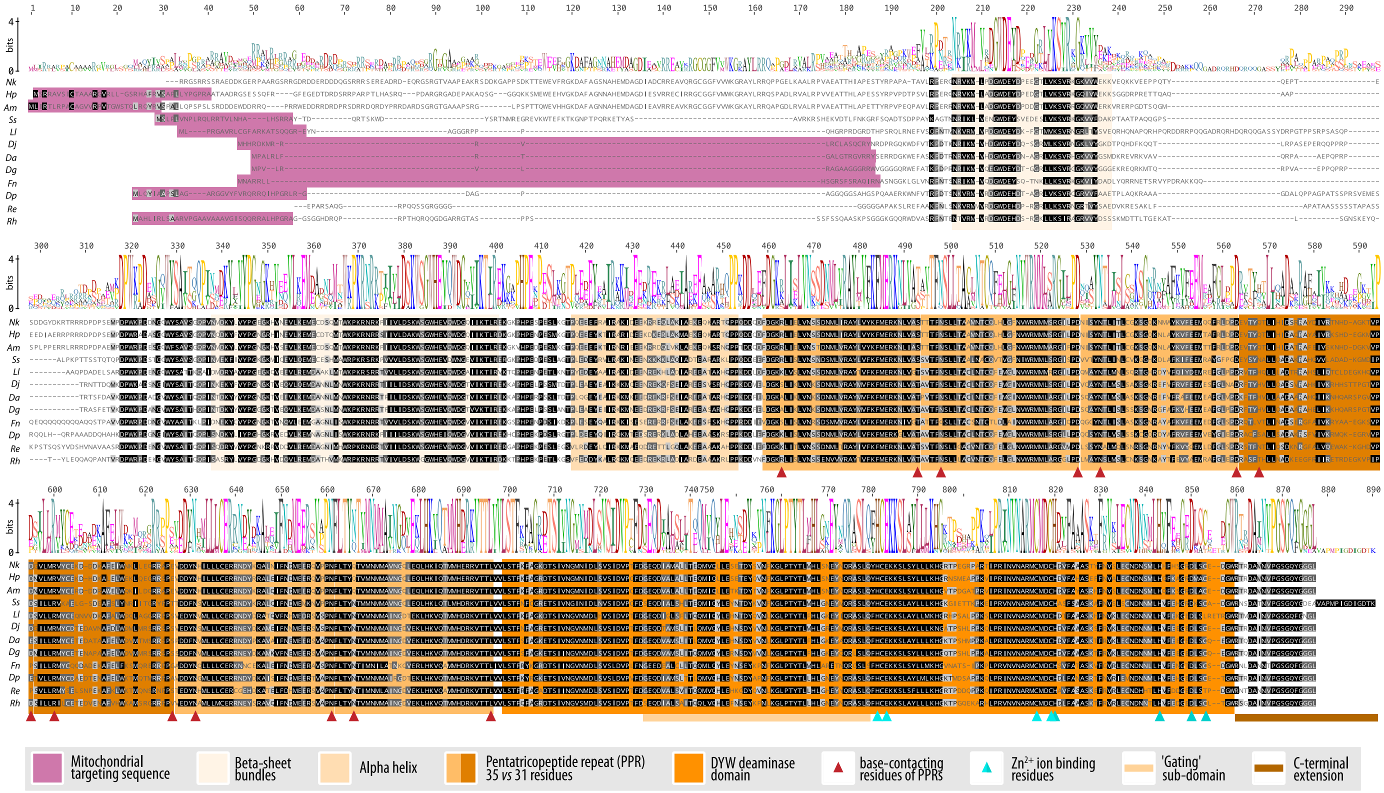


**Supplementary Figure S1**. Multiple protein-sequence alignment of PPRD1 homologs from 12 diplonemid species. Conserved sequence motifs, amino acid residues important for RNA and cofactor binding and for deamination, as well as predicted notable structural features are highlighted. Elevated residue conservation is indicated in more intense grey-scale shading. Species abbreviations: Am, *Artemidia motanka*; Da, *Diplonema ambulator*; Dg, *D. aggregatum*; Dj, *D. japonicum*; Dp, *D. papillatum*; Fn, *Flectonema neradi*; Hp, *Hemistasia phaeocysticola*; Ll, *Lacrimia lanifica*; Nk, *Namystynia karyoxenos*; Re, *Rhynchopus euleeides*; Rh, *R. humris*; Ss, *Sulcionema specki*.

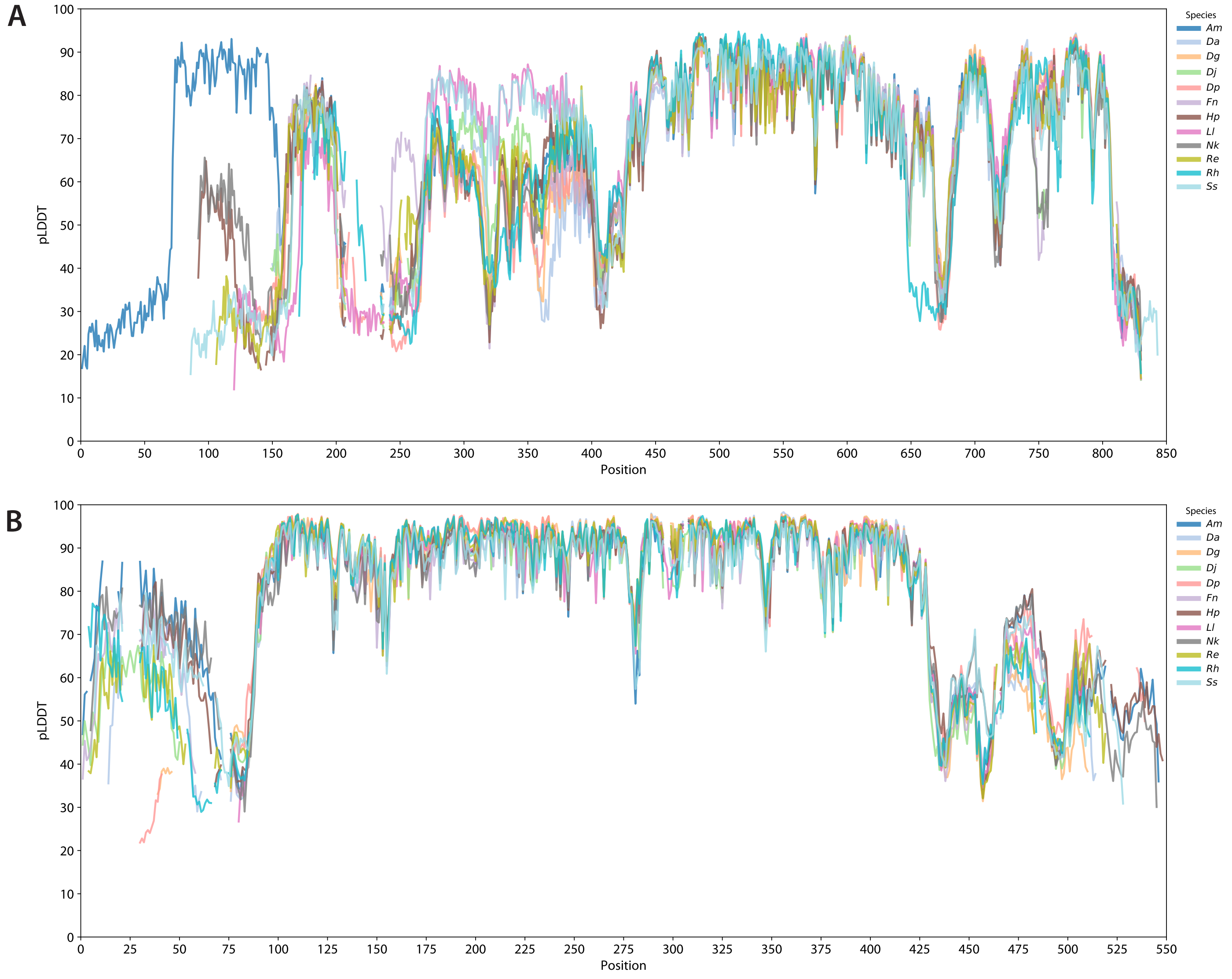


**Supplementary Figure S2**. Local confidence of AlphaFold3-predicted structural models of (**A**) PPRD1 and (**B**) DAPX1 homologs from 12 diplonemid species. pLDDT values per position in a structure-guided multiple sequence alignment; residue numbering as in **Supplementary Figure S7**. Species abbreviations: Am, *Artemidia motanka*; Da, *Diplonema ambulator*; Dg, *D. aggregatum*; Dj, *D. japonicum*; Dp, *D. papillatum*; Fn, *Flectonema neradi*; Hp, *Hemistasia phaeocysticola*; Ll, *Lacrimia lanifica*; Nk, *Namystynia karyoxenos*; Re, *Rhynchopus euleeides*; Rh, *R. humris*; Ss, *Sulcionema specki*.

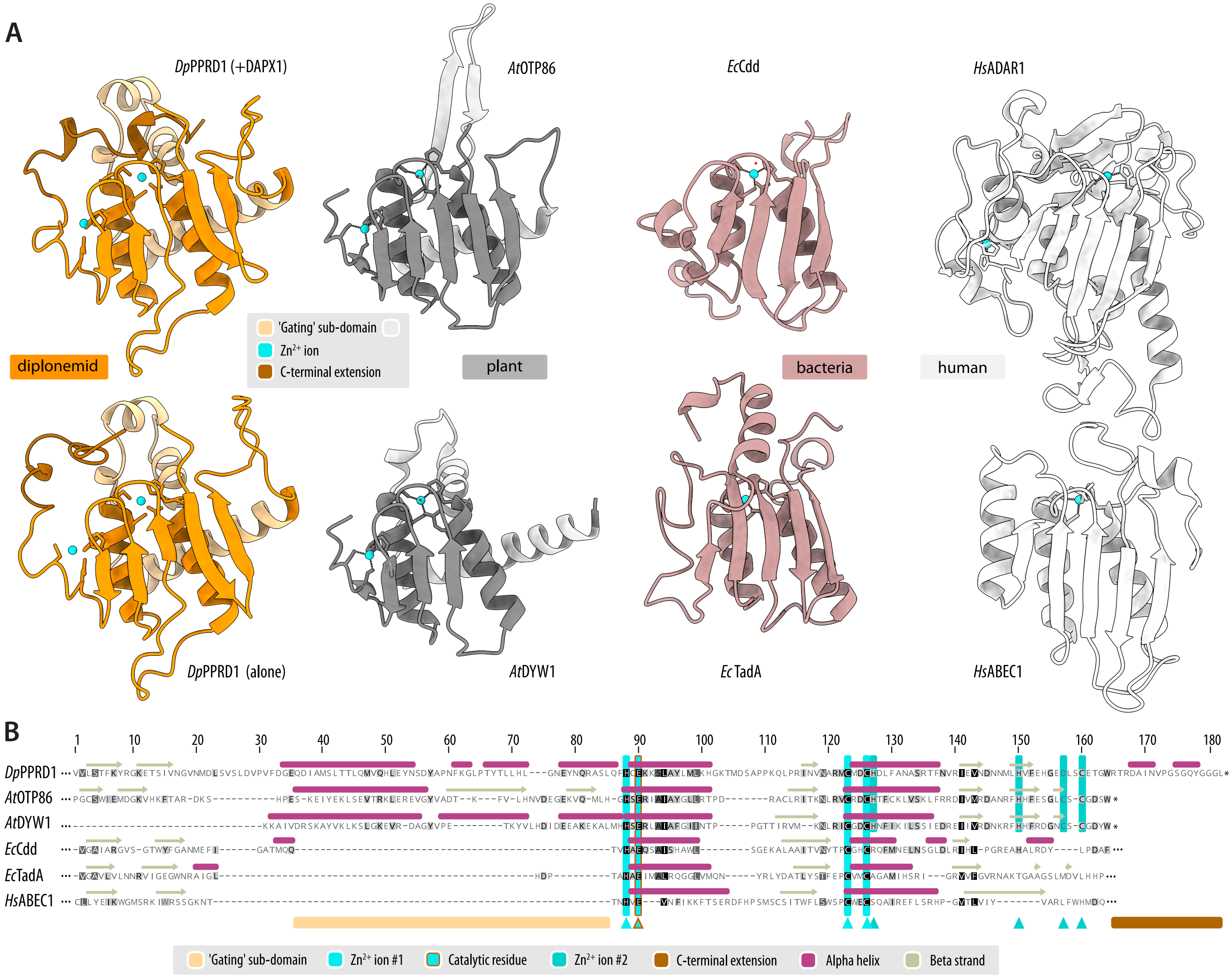


**Supplementary Figure S3**. Spatial arrangement of function-critical features in deaminase domains.
(**A**) Comparison of the predicted structures of *D. papillatum* PPRD1 (in a complex with DAPX1 [top] or alone [bottom]) to the experimentally determined structures of *Arabidopsis thaliana* DYW-family RNA deaminases OTP86 (PDB 7O4F) and DYW1 (PDB 7W86), *Escherichia coli* CMP/dCMP deaminase Cdd (PDB 1AF2), tRNA-specific adenosine deaminase TadA (PDB 8E2P), *Homo sapiens* dsRNA adenosine deaminase ADAR1 (PDB 9B84), and C-to-U-editing enzyme APOBEC-1/ABEC1 (PDB 6X91). The N-terminus of the DYW domain represented by its first two beta strands is often referred to as the 'PG box' named after the first two Pro-Gly residues, which sequence-wise, are moderately conserved across most plant sequences, especially those that carry PPR arrays [1]. Note that *At*DYW1 lacks the two beta strands and thus the 'PG box' (see also (**B**)). Active site residue and zinc-ion binding residues are represented as sticks. The 'gating' sub-domain is highlighted in lighter tints, diplonemid C-terminal extension in darker shades, and $Zn^{2+}$ ions in cyan. Note that only the DYW-family proteins include the second zinc ion-binding site.
(**B**) Annotated structure-guided multiple sequence alignment of proteins carrying the cytidine-deaminase fold shown in (**A**), namely *Dp*PPRD1 (DIPPA_21441), *At*OTP86 (Uniprot ID Q9M1V3), *At*DYW1 (P0C7R1), *Ec*Cdd (P0ABF6), *Ec*TadA (P68398), *Hs*ABEC1 (P41238). Despite certain structural similarities, that *Hs*ADAR1 (P55265), a member of the dsRNA adenosine deaminase family, is not included because this family has a distinctive catalytic site architecture and a different distribution of active site residues along the sequence compared to the cytidine deaminase superfamily.

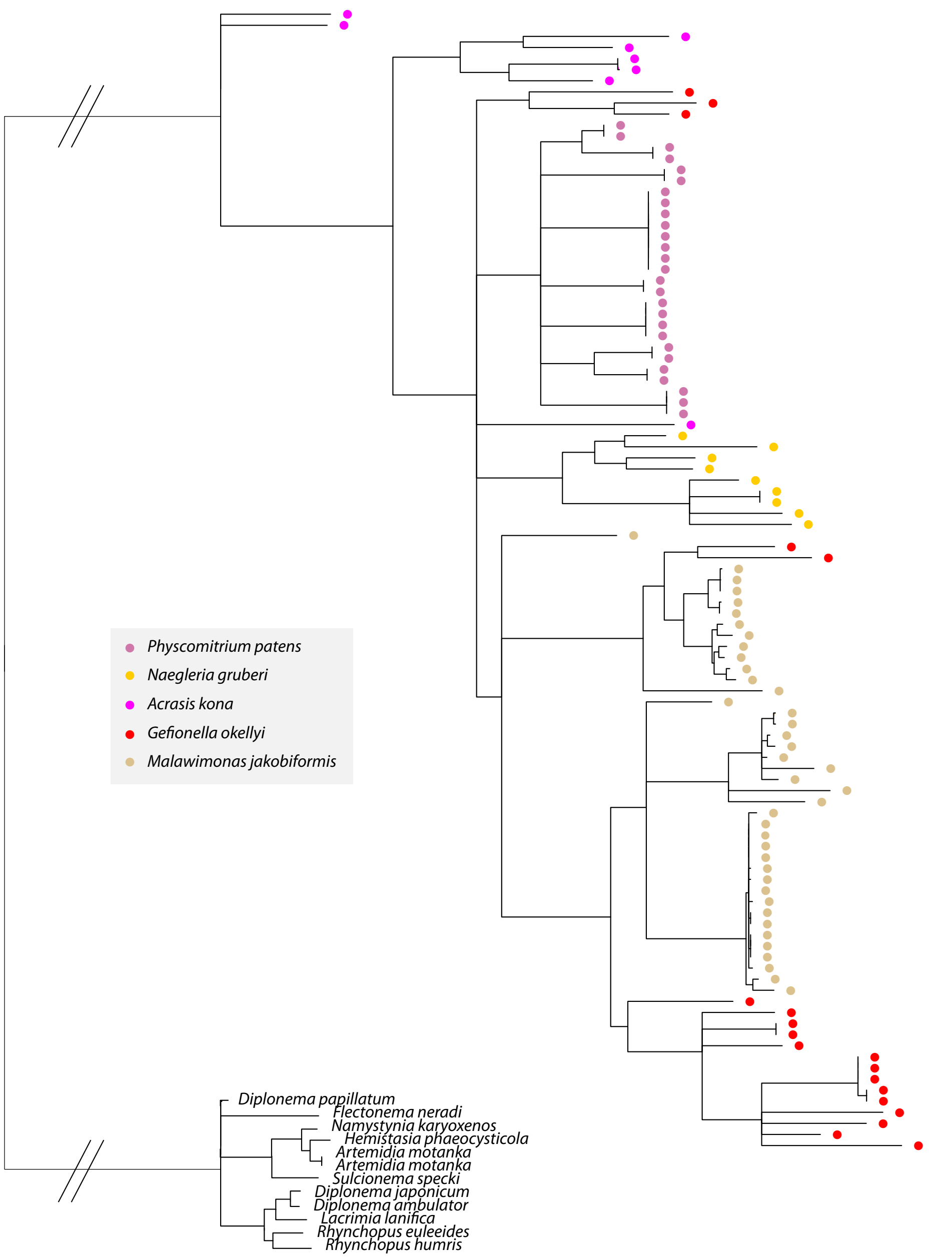


**Supplementary Figure S4**. Phylogenetic analysis of the DYW domain from Discoba, Malawimonada and *Physcomitrium*. Only alignment positions assigned a posterior probability of 1 were retained, yielding 172 amino acid positions for phylogenetic analysis. Closely related sequences such as the diplonemid proteins and species-specific paralog groups generally form well-supported clusters, but due to the small dataset, deeper relationships among taxa remain poorly resolved. Most paralogs from non-diplonemid taxa cluster by species, consistent with independent expansions of the PPR-DYW family. Several malawimonad homologs group across species boundaries, suggesting that multiple copies were already present in their common ancestor and subsequently differentially retained or expanded.

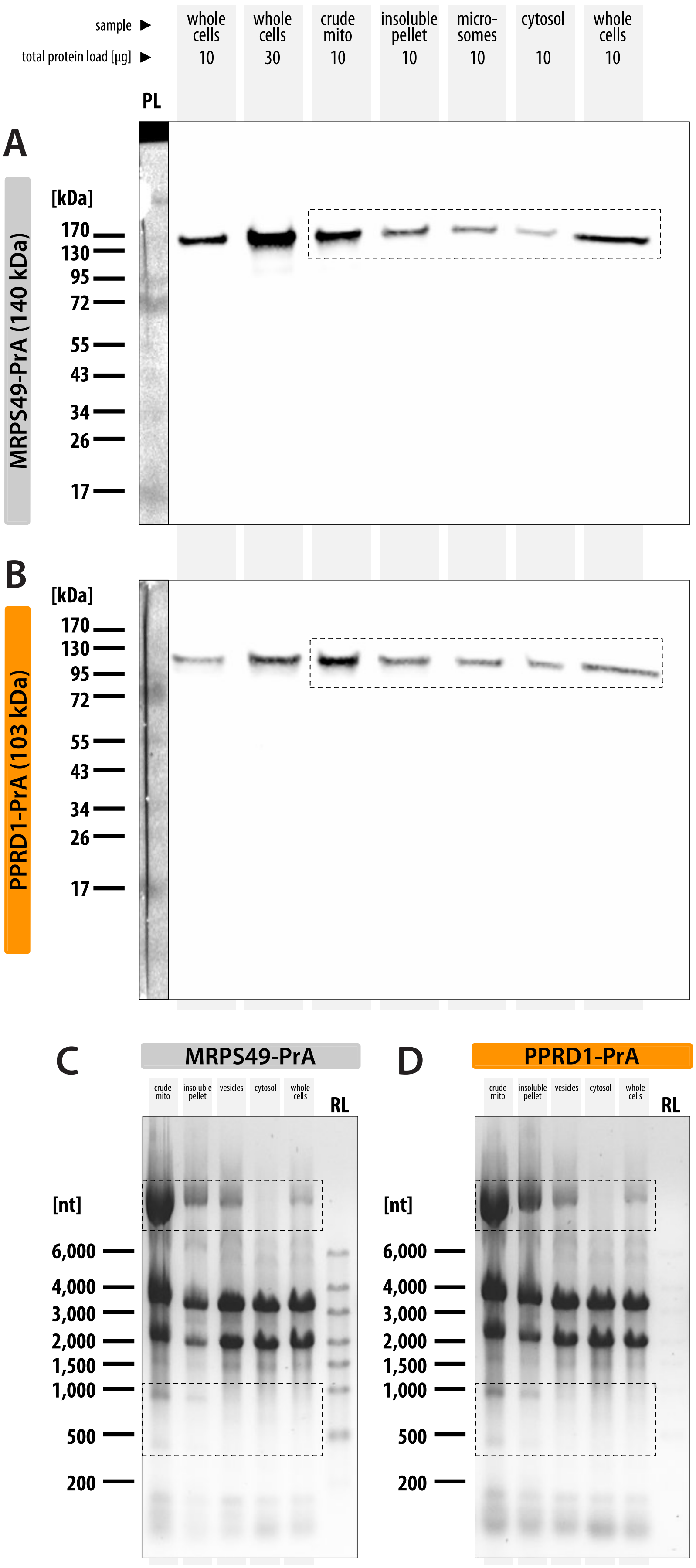


**Supplementary Figure S5**. *Diplonema* sub-cellular fractionation.

(**A**, **B**) Western blot of fractions from cell lines expressing PrA-tagged mitoribosomal protein MRPS49 (**A**) and PrA-tagged PPRD1 (**B**) separated on a discontinuous sucrose velocity gradient. Collected fractions: cytosol (solute above the top sucrose layer of 36%), microsomes (interface between the top layer and 36% sucrose), insoluble pellet (pellet below the 60% sucrose layer), crude mitochondria (interface between the 36% and 60% sucrose). Whole cell lysates were used as baseline protein-level references. PL, protein ladder. Dashed boxes indicate areas shown in **Figure 1**.

(**C**, **D**) Ethidium-bromide-stained agarose gels of nucleic acids in fractions of a discontinuous sucrose velocity gradient, obtained from cell lines expressing PrA-tagged mito-ribosomal protein MRPS49 (**C**) and PrA-tagged PPRD1 (**D**). RL, RNA ladder. Dashed boxes indicate areas shown in **Figure 1**: the box at the top delineates mitochondrial DNA, while the bottom box delimits the area containing the large and small mitoribosomal RNAs.

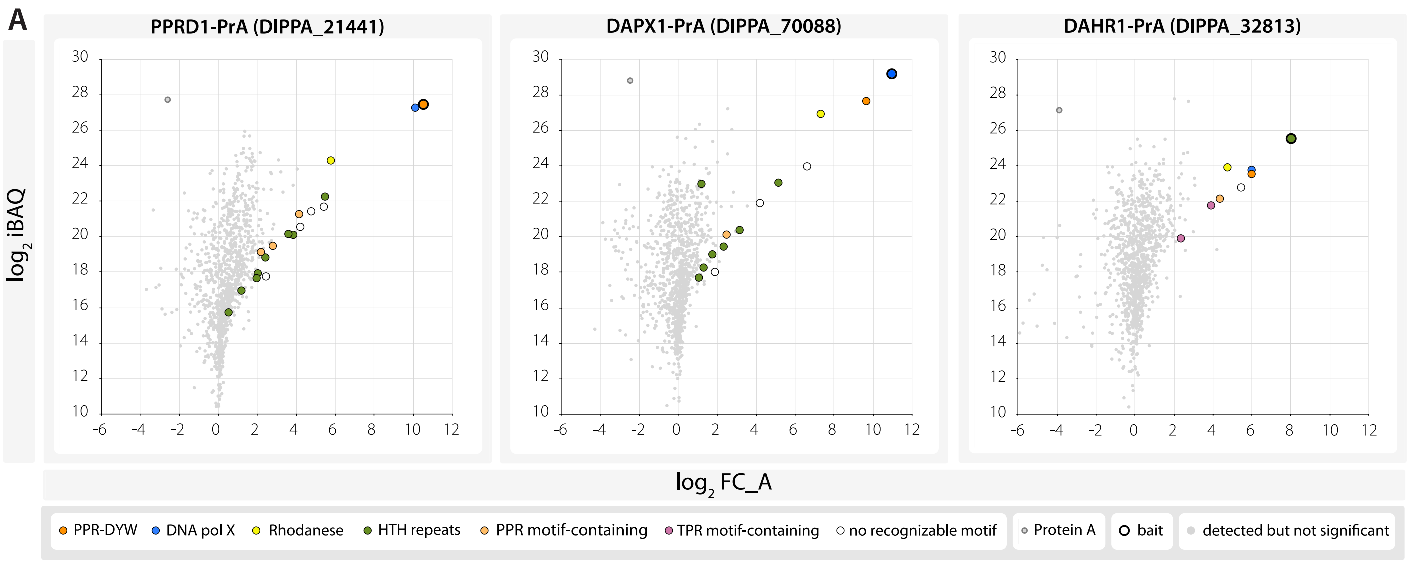


**Supplementary Figure S6**. Abundance-vs-enrichment plots of proteins pulled down together with PrA-tagged PPRD1 (DIPPA_21441, left panel), DAPX1 (DIPPA_70088, middle), and DAHR1 (DIPPA_32813, right). Proteins were classified based on their enrichment (average fold change; FC_A), abundance (iBAQ), and probability score (SP > 0.9) (for details, see **Supplementary Table S5**). Only significant proteins were highlighted in different colours based on the presence of characteristic domains and/or sequence motifs (e.g., DNA PolX, PPR, TPR).

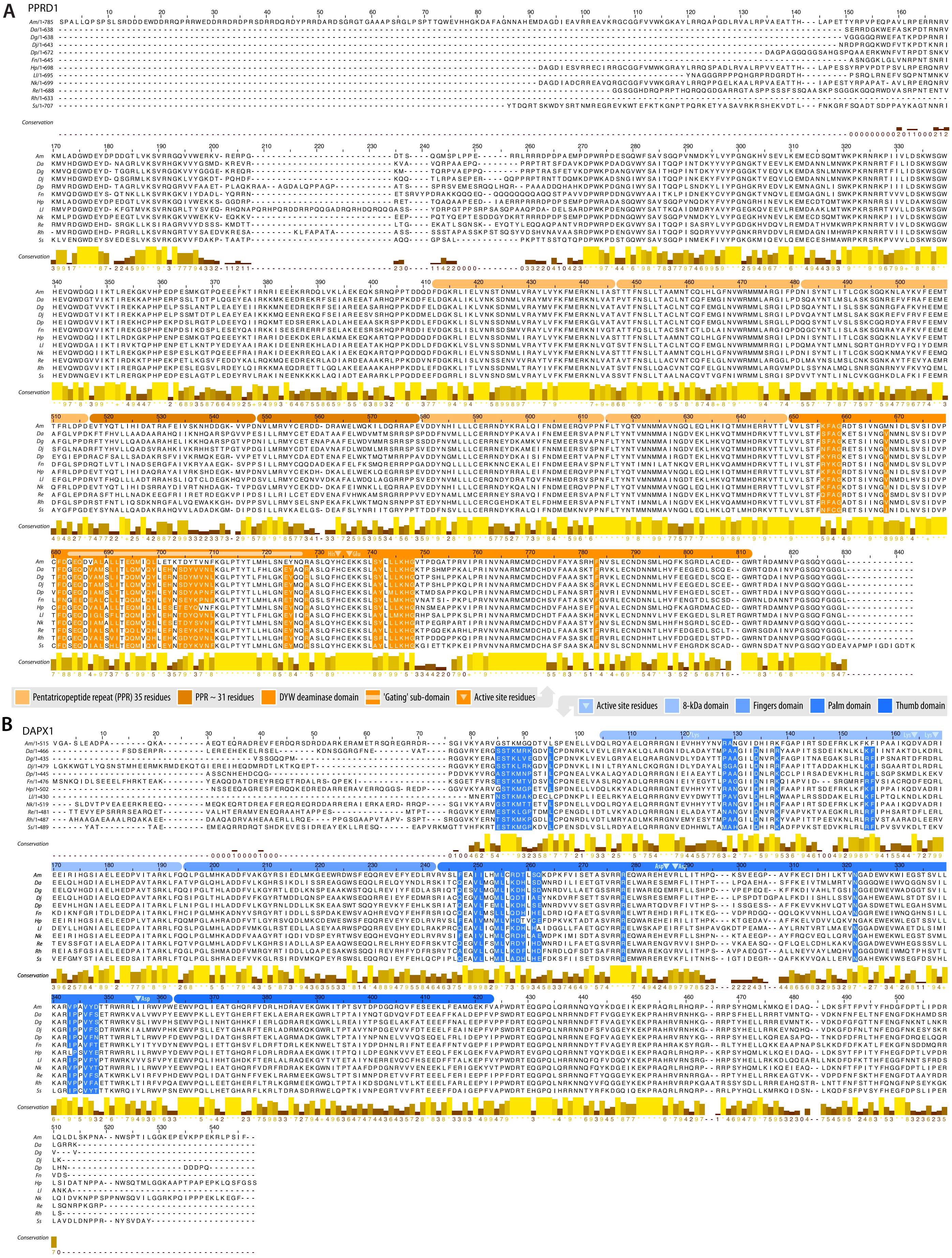
A
PPRD1
B
DAPX1
Conservation
Pentatricopeptide repeat (PPR) 35 residues
PPR ~ 31 residues
DYW deaminase domain
'Gating' sub-domain
Active site residues
Active site residues
8-kDa domain
Fingers domain
Palm domain
Thumb domain

**Supplementary Figure S7**. Structure-guided multiple sequence alignment of (**A**) PPRD1 and (**B**) DAPX1 homologs from 12 diplonemid species with predicted conserved interacting residues highlighted in orange and blue, respectively. Domains and other salient features of the two proteins are highlighted in the scale bar, as described in the keys. Also indicated are identities of conserved active site residues expected in a catalytically active enzyme. Species abbreviations: Am, *Artemidia motanka*; Da, *Diplonema ambulator*; Dg, *D. aggregatum*; Dj, *D. japonicum*; Dp, *D. papillatum*; Fn, *Flectonema neradi*; Hp, *Hemistasia phaeocysticola*; Ll, *Lacrimia lanifica*; Nk, *Namystynia karyoxenos*; Re, *Rhynchopus euleeides*; Rh, *R. humris*; Ss, *Sulcionema specki*.

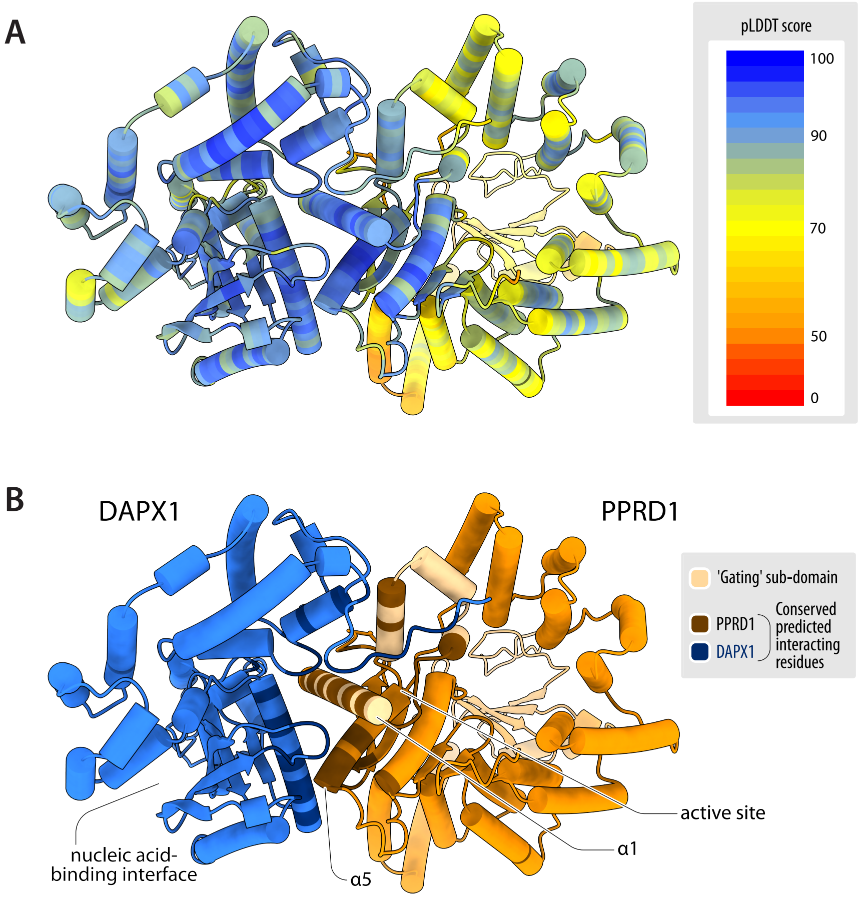


**Supplementary Figure S8**. Predicted structure of the PPDR1–DAPX1 binary complex.
(**A**) Local confidence of AlphaFold3-predicted structural models of PPRD1 and DAPX1 in the PPRD1–DAPX1 binary complex. pLDDT values per position are coloured as shown in the key. Score interpretation: 90–100, high expected accuracy; 70–90, high confidence for the model backbone but sides chains may be inaccurate; 50–70, low confidence; <50, very low confidence, possibly disordered or flexible [12].
(**B**) Interacting residues (shown in darker shades; **Supplementary Figure S7**) were inferred from residue pair distances (≤4 Å) in the complex structure, conserved across diplonemids in regions with reliable local structure. Low-confidence regions of pLDDT<50 at N- and C-termini are not shown.

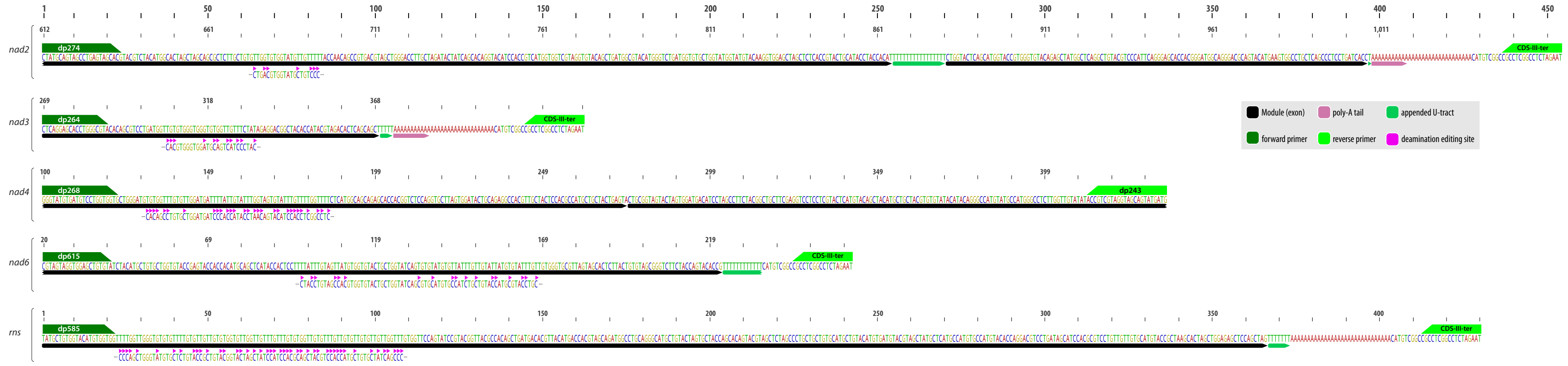


**Supplementary Figure S9**. RT-PCR amplicons generated to analyze the influence of gene silencing on *in vivo* deamination RNA editing. The depicted sequences represent the prevalent mature mRNAs as determined previously [24,76], together with annotations of modules (transcript pieces), and sites of uridylation-appendage editing, terminal polyadenylation, and deamination editing. Clusters of deamination RNA editing sites, with the pre-edited sequences shown below that of the mature mRNA, are located in the penultimate module of *nad2* (‘m4’), in the last module of *nad3* (‘m2’), and in the first modules of *nad4*, *nad6*, and *rns* (‘m1’). Reverse transcription (RT) was done using an oligonucleotide complementary to the mRNA’s native poly-A tail, except for i) *nad6*, in which case the oligo was complementary to its internal 50 U-long tract, and ii) *rns*, which was artificially poly-adenylated prior to RT. The topmost ruler indicates positions along the amplicon, while the ruler above each of the five amplicons indicates the position along the corresponding mature mRNA. Note that *nad2*, *nad3*, and *nad6* were initially referred to as *y3*, *y1*, and *y5* [29].
